# Aggressive mimicry lure polymorphisms in the parasitic mussel *Lampsilis fasciola* model fish or leech host prey and differ in morphology and pigmentation, but not in display behavior

**DOI:** 10.1101/2023.11.27.568842

**Authors:** Trevor L. Hewitt, Paul Johnson, Michael Buntin, Talia Moore, Diarmaid Ó Foighil

## Abstract

Unionoid freshwater mussels (Bivalvia: Unionidae) are free-living apart from a brief, obligately parasitic, larval stage that infects fish hosts and gravid female mussels have evolved a spectrum of strategies to infect fish hosts with their larvae. In many North American species, this involves displaying a mantle lure: a pigmented fleshy extension that acts as an aggressive mimic of a host fish prey, thereby eliciting a feeding response that results in host infection. The mantle lure of *Lampsilis fasciola* is of particular interest because it is apparently polymorphic, with two distinct primary lure phenotypes. One, described as “darter-like”, has “eyespots”, a mottled body coloration, prominent marginal extensions, and a distinct “tail”. The other, described as “worm-like”, lacks those features and has an orange and black coloration. We investigated this phenomenon to 1) confirm that it is a true polymorphism; 2) investigate its ecological persistence; 3) identify the range of putative model species targeted by this mimicry system within a river drainage; 4) determine whether the mantle lure polymorphism includes a behavioral component. Detection of within-brood lure variation and within-population phylogenomic (ddRAD-seq) analyses of individuals bearing different lures confirmed that this phenomenon is a true polymorphism. It appears stable over ecological timeframes: the ratio of the two lure phenotypes in a River Raisin (MI) population in 2017 was consistent with that of museum samples collected at the same site 6 decades earlier. Within the River Raisin, four main “darter-like” lure motifs visually approximated four co-occurring darter species (*Etheostoma blennioides, E. exile, E. microperca,* and *Percina maculata*) and the “worm-like” lure resembled a widespread common leech, *Macrobdella decora*. Darters and leeches are typical prey of *Micropterus dolomieui* (smallmouth bass), the primary fish host of *L. fasciola*. *In situ* field recordings were made of the *L. fasciola* “darter” and “leech” lure display behaviors, in addition to the non-polymorphic lure display of co-occurring *L. cardium*. Despite having putative models in distinct phyla, both *L. fasciola* lure morphs have similar display behaviors that differ significantly from that of sympatric *L. cardium* individuals. We conclude that the *L. fasciola* mantle lure polymorphism does not include a behavioral component. Discovery of discrete within-brood inheritance of the lure polymorphism implies potential control by a single genetic locus and identifies *L. fasciola* as a promising study system to identify regulatory genes controlling a key adaptive trait of freshwater mussels.

## INTRODUCTION

In ecology, mimicry refers to a convergent adaptive trait prevalent in many biological communities: the deceptive resemblance of one organism to another (Pasteur, 1982; Schaefer & Ruxton, 2009; Maran, 2015). It involves three categories of interacting ecological players: mimic (organism displaying the deceptive resemblance), model (organism being mimicked), and receiver (organism being deceived) (Pasteur, 1982; Maran, 2015). Mimicry occurs across a wide variety of ecological contexts and sensory modalities, but conceptually (Jamie, 2017), individual cases can be categorized by the traits being mimicked (signals or cues), as well as by the degree of deceptiveness and the fitness consequences being communicated to the receiver (aggressive, rewarding, Müllerian or Batesian mimicry).

Mimetic systems that are polymorphic (multiple within-species mimic morphs with discrete models) have been particularly influential in uncovering the genetic basis of complex adaptive traits in natural populations (Jay et al., 2018; Palmer & Kronforst, 2020). Such polymorphisms are rare in nature, with the most well studied examples occurring in papilionid butterflies (Hazel, 1990; Joron & Mallet, 1998; Nijhout, 2003; Jay et al., 2018; Palmer & Kronforst, 2020). For instance, polymorphisms in *Heliconious* species are determined by presence/absence of an introgressed chromosomal inversion ‘supergene’ (Jay et al., 2018), and alleles of a single ancestral gene (*doublesex*) control female-specific polymorphisms in *Papilio* species (Palmer & Kronforst, 2020).

In contrast to papilionid butterflies, the genetics of mimicry trait evolution among unionoid mussels is poorly understood. Unionoida comprise ∼75% of the planet’s freshwater bivalve species and are free-living apart from a brief, obligately parasitic, larval stage that infects fish hosts (Bogan, 2007; Haag, 2012). Gravid female mussels have evolved a spectrum of strategies to infect hosts with their larvae (Zanatta & Murphy, 2006; Barnhart, Haag & Roston, 2008; Hewitt, Wood & Ó Foighil, 2019). Females in many species use a mantle lure (Welsh, 1933): a pigmented fleshy extension that acts as an aggressive mimic (Jamie, 2017) of a host fish prey item (e.g., small fish, aquatic invertebrate, egg mass) visual cue and elicits a feeding response resulting in host infection (Haag & Warren, 1999; Barnhart, Haag & Roston, 2008; Figure 1a). Mimetic mantle lures predominate in Lampsilini, a major clade of North American freshwater mussels recently identified as a cryptic adaptive radiation centered on larval ecologies and specialized host-infection behaviors (Hewitt, Haponski & Ó Foighil, 2021). Ortmann (1911) and Kramer (1970) reported the production of rudimentary mantle lures in juveniles and male lampsilines but noted that formation of fully developed lures is restricted to sexually mature females, and that only gravid females engage in lure display behaviors. Surprisingly, neither Ortmann (1911) nor Kramer (1970) depicted male mussel lure rudiments, nor could we find any such depictions in the literature, and we therefore included examples for two species, *Lampsilis cardium* and *L. fasciola* (Supplementary Figure 1).

**Figure 1:**
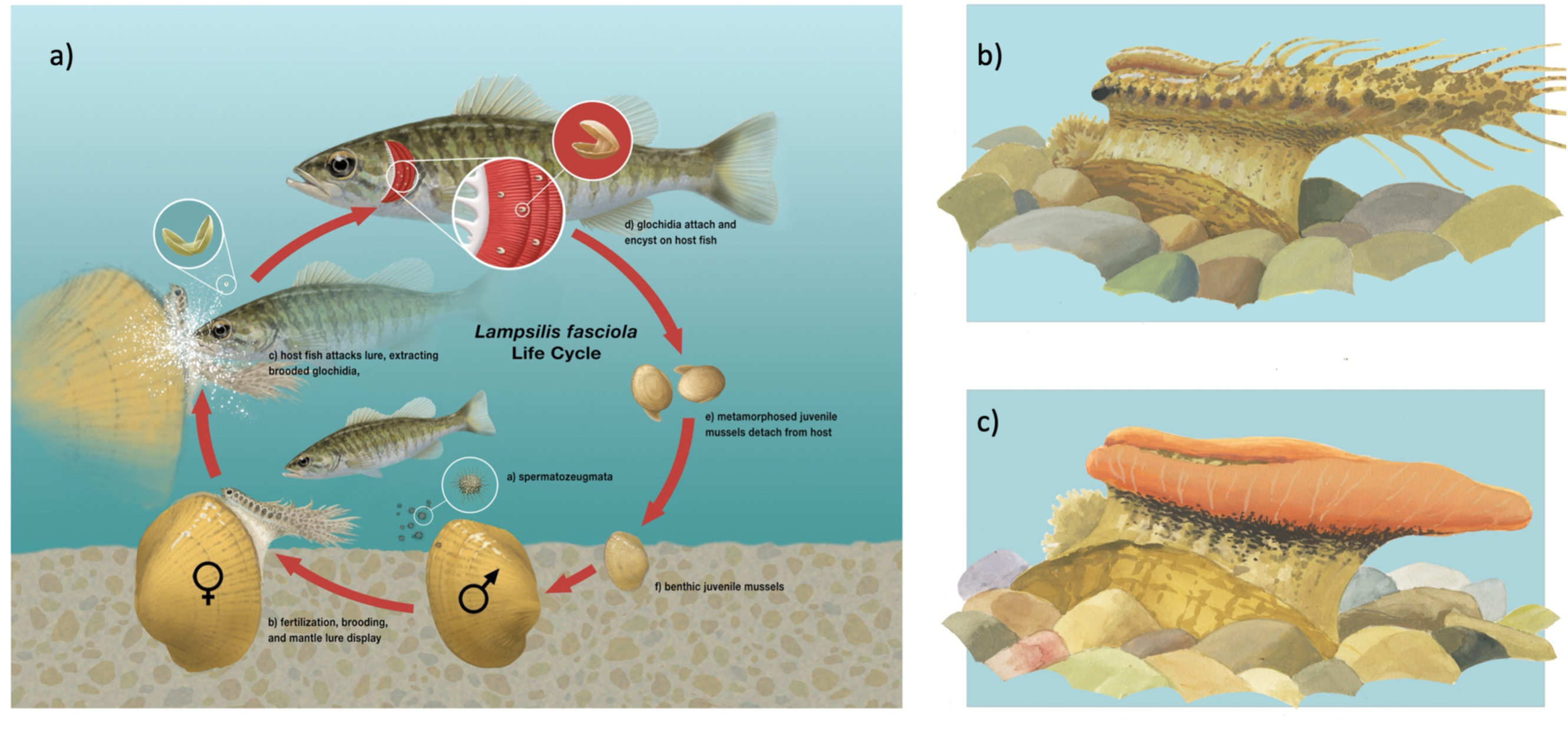
a) Summary diagram representing the life cycle of the freshwater mussel *Lampsilis fasciola.* After fertilization, the gravid female mussel displays a mantle lure, here a darter mimic, to the primary fish host, *Micropterus dolomieu*. This elicits an attack through which the host is infected by mussel parasitic larvae (glochidia). After a short infective period (∼2 weeks), the parasitic larvae metamorphose into juvenile mussels that detach from the host and fall to the sediment. Panels b (“darter-like”) and c (“worm-like”) depict the two primary phenotypes of lure observed in *L. fasciola*. The former (b) has “eyespots”, a mottled “main body” pigmentation composed of lateral and dorsal spots that can vary substantially in color, numerous and prominent marginal extensions, and a distinct “tail” region, whereas the latter lacks those features and has instead a uniform bright orange coloration underlain with a black basal stripe

Although mimetic mantle lures are a key adaptive trait of freshwater mussel diversification, the genetic regulators underlying their formation (Kramer, 1970), variation (Haag, Warren & Shillingsford, 1999; Zanatta, Fraley & Murphy, 2007; Barnhart, Haag & Roston, 2008), and evolution (Zanatta & Murphy, 2006; Hewitt, Haponski & Ó Foighil, 2021) remain completely unknown. This gap in our knowledge is exacerbated by the stark conservation status of North American freshwater mussels, with 2/3rds of species classified as threatened or near-threatened (Lopes-Lima et al., 2018). As with papilionid butterflies (Jay et al., 2018; Palmer & Kronforst, 2020), targeting polymorphic lampsiline mantle lures for in-depth study may represent a tractable route to closing that gap between genes and phenotypes.

*Lampsilis fasciola,* the Wavy-Rayed Lampmussel, is a promising candidate species in that it produces a number of distinct mantle lure phenotypes (Zanatta, Fraley & Murphy, 2007) across its Eastern North America distribution, extending from southern Ontario to northern Alabama (Parmalee & Bogan, 1998). Two range-wide lure phenotypes predominate in northern populations. The more common of the two, labeled “darter-like” by Zanatta et al. (2007), has “eyespots”, a mottled “main body” pigmentation composed of lateral and dorsal spots that can vary substantially in color, numerous and prominent marginal extensions (AKA “appendages” or “tentacles”), and a distinct “tail” region (Kramer, 1970; Zanatta, Fraley & Murphy, 2007) – see Figure 1b. A rarer lure phenotype, labeled “worm-like” by McNichols (2007), lacks the above features and has instead a uniform bright orange coloration underlain with a black basal stripe (Zanatta, Fraley & Murphy, 2007) – see Figure 1c. The latter lure phenotype is highly distinctive within the genus *Lampsilis* where fish-like mantle lures are the norm (Kramer, 1970). Based on the results of laboratory larval infection experiments, and on the degree of ecological overlap, *Micropterus dolomieu* (Smallmouth Bass), and to a lesser extent *Micropterus salmoides* (Largemouth Bass), have been identified as *Lampsilis fasciola’s* primary fish hosts (Zale & Neves, 1982; McNichols, 2007; Morris et al., 2009; McNichols, Mackie & Ackerman, 2011; VanTassel et al., 2021). Both host species are generalist predators of aquatic invertebrates and vertebrates (Clady, 1974).

Our study aimed to address outstanding, inter-related questions to develop *Lampsilis fasciola* into an integrated mantle lure polymorphism study system. First among them was residual uncertainty that the mantle lure morphs represent polymorphisms rather than cryptic species. Zanatta et al. (2007), using microsatellite markers, did not detect evidence of cryptic species but qualified their conclusions due to small sample sizes, and their result require molecular phylogentic corroboration (Fisheries and Oceans Canada, 2018). Secondly, we lack any data on the relative persistence of the lure polymorphisms in natural populations through time. Thirdly, one set of important ecological players – the respective models of each *L. fasciola* mantle lure mimic – has not been determined with any specificity. Finally, mantle lure display behavior is an important component of effective mimicry in freshwater mussels (Welsh, 1933; Jansen, Bauer & Zahner-Meike, 2001; Haag & Warren, 2003; Barnhart, Haag & Roston, 2008), but it is unknown if morphologically divergent *L. fasciola* mantle lures, that presumably mimic very distinct host prey models, also differ in their display behaviors.

We leveraged a combination of field-collection, captive breeding, museum specimens, and ecological surveys to collect genetic, phenotypic, and population data on this species. Using novel data from two southeastern Michigan river populations [the Raisin (our primary study location), and the Huron], and a phylogenomic (ddRADseq) approach, we corroborated Zanatta et al.’s (2007) finding that mussels bearing the two mantle lure morphs do not represent cryptic lineages. This was confirmed by our documentation of discrete, within-brood, inheritance of the *L. fasciola* lure polymorphism in the captive specimens from the Alabama Aquatic Biodiversity Center (AABC). Availability of mid-20th century museum specimens from a River Raisin population revealed that the relative frequency of the two morphs has remained broadly stable over 60 years. In addition, availability of a comprehensive Raisin River ichtyofauna survey (Smith, Taylor & Grimshaw, 1981) allowed us to identify four darter species as putative models for the predominant “darter-like” mantle lures in this population. The most palusible model for the “worm-like” *L. fasciola* mantle lure appears to be *Macrobdella decorata* (the American medical leech). Despite having putative models in distinct phyla, detailed video analyses of gravid females in the two drainages revealed that both *L. fasciola* lure morphs have similar display behaviors that differ significantly from that of sympatric *L. cardium* indivuals bearing non-polymorphic fish-mimic lures. We conclude that the *L. fasciola* mantle lure polymorphism does not include a significant behavioral component.

## MATERIALS AND METHODS

### Tissue Sample Collection

*Lampsilis fasciola* mantle tissue samples were collected for genotyping purposes by taking non-lethal mantle clip biopsies (Berg et al., 1995) from wild population lure-displaying female mussels during the summers of 2017, 2018, and 2021 from a total of three rivers (Figure 2). Two of the sampling locations were in southeastern Michigan: the River Raisin at Sharon Mills County Park (42.176723, -84.092453; N=30; 24 “darter-like”, 6 “worm-like”, collectively sampled in 2017, 2018 & 2020), and the Huron River at Hudson Mills Metropark, MI (42.37552, -83.91650; N=13; 7 “darter-like”, 6 “worm-like”, collectively sampled in 2017, 2018, and 2020 under the MI Threatened and endangered species collection permit number 149). Both rivers flow into Lake Erie and are part of the Saint Lawrence drainage. The third location was in North Carolina: the Little Tennessee River (N=10; 35.32324, -83.52275; N=1-, all 10 were “darter-like” and sampled in 2017); this river is a tributary of the Tennessee River and part of the Mississippi drainage. Prior to each biopsy, photographs of the intact, undisturbed, lure display were taken with an Olympus Tough TG-6 underwater camera (Supplementary figure 2).

**Figure 2:**
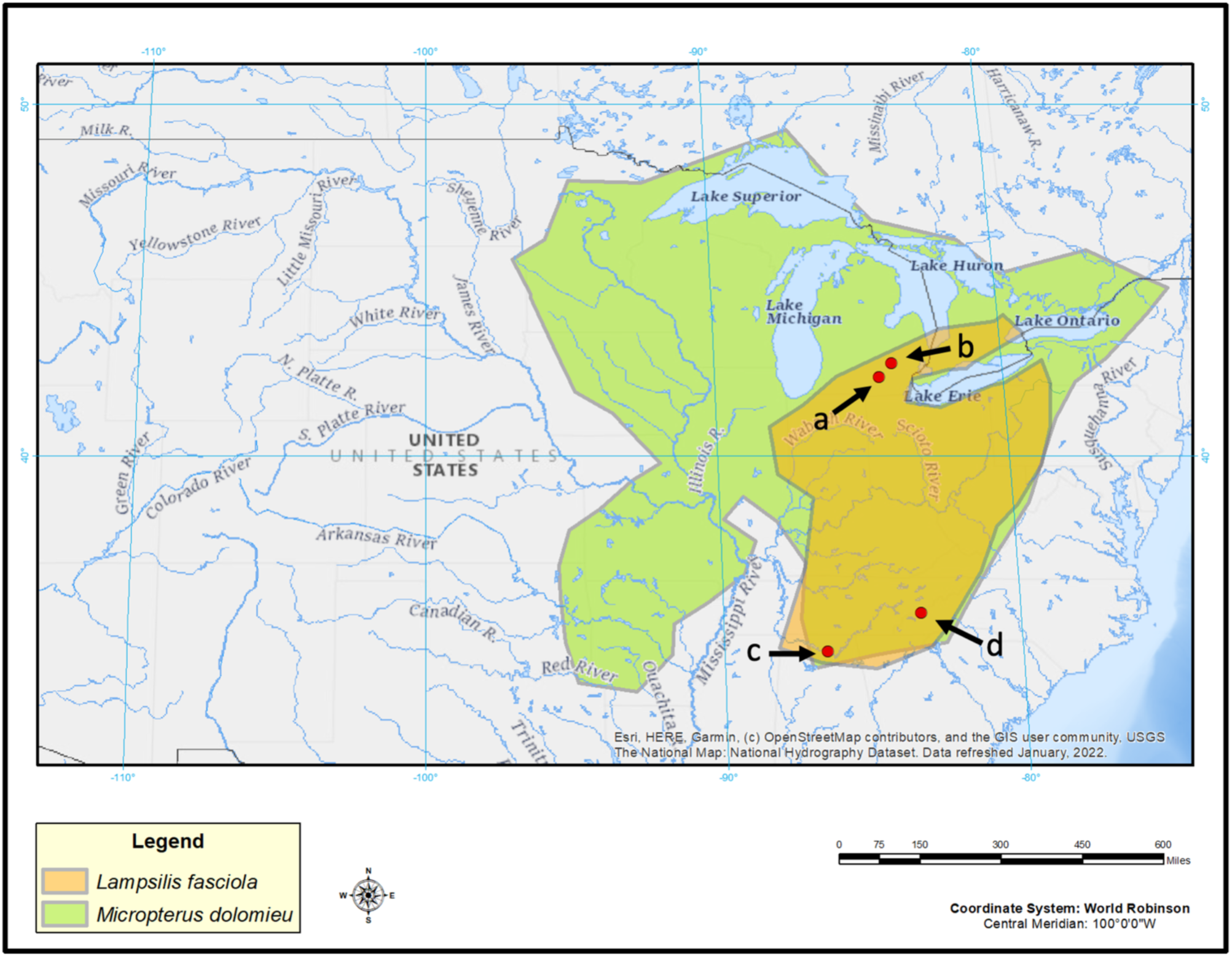
Map of Eastern North America showing the sampling sites of *Lampsilis fasciola* and the estimated ranges of this freshwater mussel species and of its primary host fish *Micropterus dolomieu*. A indicates the Raisin River sampling site at Sharon Mills County Park and B the Huron River sampling site at Hudson Mills Park, both in southeastern Michigan. Site C indicates the Paint Rock River sampling site in Alabama, and site D is the Little Tennessee River site in North Carolina.

### Captive Brood Tissue Samples

In addition to the wild-sampled specimens, we also obtained tissue samples from 50 members of a captive-raised individuals of a single brood that had been ethanol-preserved a decade earlier. In 2009, the Alabama Aquatic Biodiversity Center (AABC) established a culture facility for endangered freshwater mussels. The Center’s inaugural culture attempt, by co-authors Paul Johnson and Michael Buntin, was a proof-of-concept trial involving a single gravid female *Lampsilis fasciola*, a non-endangered species, sourced from a wild Paint Rock River (another Tennessee River tributary) population (N 34˚ 47.733′,W 86˚ 14.396′) in Jackson County, AL (Figure 2) on June 11, 2009. This female *L. fasciola* had a “worm-like” lure: the AABC data sheet for the trial 2009 host infection (Supplemental Figure 3) records that it was “bright orange and black”, and it lacked the “eyespots”, mottled body coloration, marginal extensions, and “tail” of the “darter-like” lure phenotype (Butlin & Johnson, pers. observ.). On July 13^th^ 2009, ∼31K glochidia larvae were extracted from the female’s marsupia and used to infect *Micropterus coosae* (Redeye Bass) hosts sourced from the Eastaboga Fish Hatchery (Calhoun County, AL) using standard protocols (Barnhart, Haag & Roston, 2008). The female mussel was then returned live to the Paint Rock River Population. Following completion of larval development on the fish hosts, ∼ 9.3K metamorphosed juvenile mussels were recovered and reared, initially for the first few weeks in mucket bucket systems (Barnhart, 2006) then a suspended upwelling system (SUPSYS) for two years with ∼2.2K surviving. In 2011, this proof-of-concept culture experiment was terminated, and the survivors were donated to several research groups, with the large majority utilized for toxicology experiments (Leonard et al., 2014a,b).

Prior to the brood’s termination, Johnson noticed that a few females had attained sexual maturity and were displaying polymorphic lures (Figures 3b, 3c). We aimed to substantiate that 2011 observation by tracking down any remaining brood specimens and were successful in recovering 50 individuals that had been preserved in 95% ethanol and shipped to Nathan Johnson (USGS) in Gainsville, FL in 2011. Because *Lampsilis* spp. juveniles and males produce a rudimentary mantle lure (Ortmann, 1911; Kramer, 1970), we were able to determine the primary lure phenotype (darter-like” or “worm-like”) of all 50 preserved brood members. Using a Leica MZ16 dissecting microscope, individual photomicrographs were taken of the preserved rudimentary lure structures (Figure 3d,e and Supplementary Figure 4), and their respective lure phenotypes were identified independently by both T. Hewitt and by D. Ó Foighil. Additionally, tissue samples were acquired from all 50 individuals and included in the downstream phylogenomic analyses.

**Figure 3:**
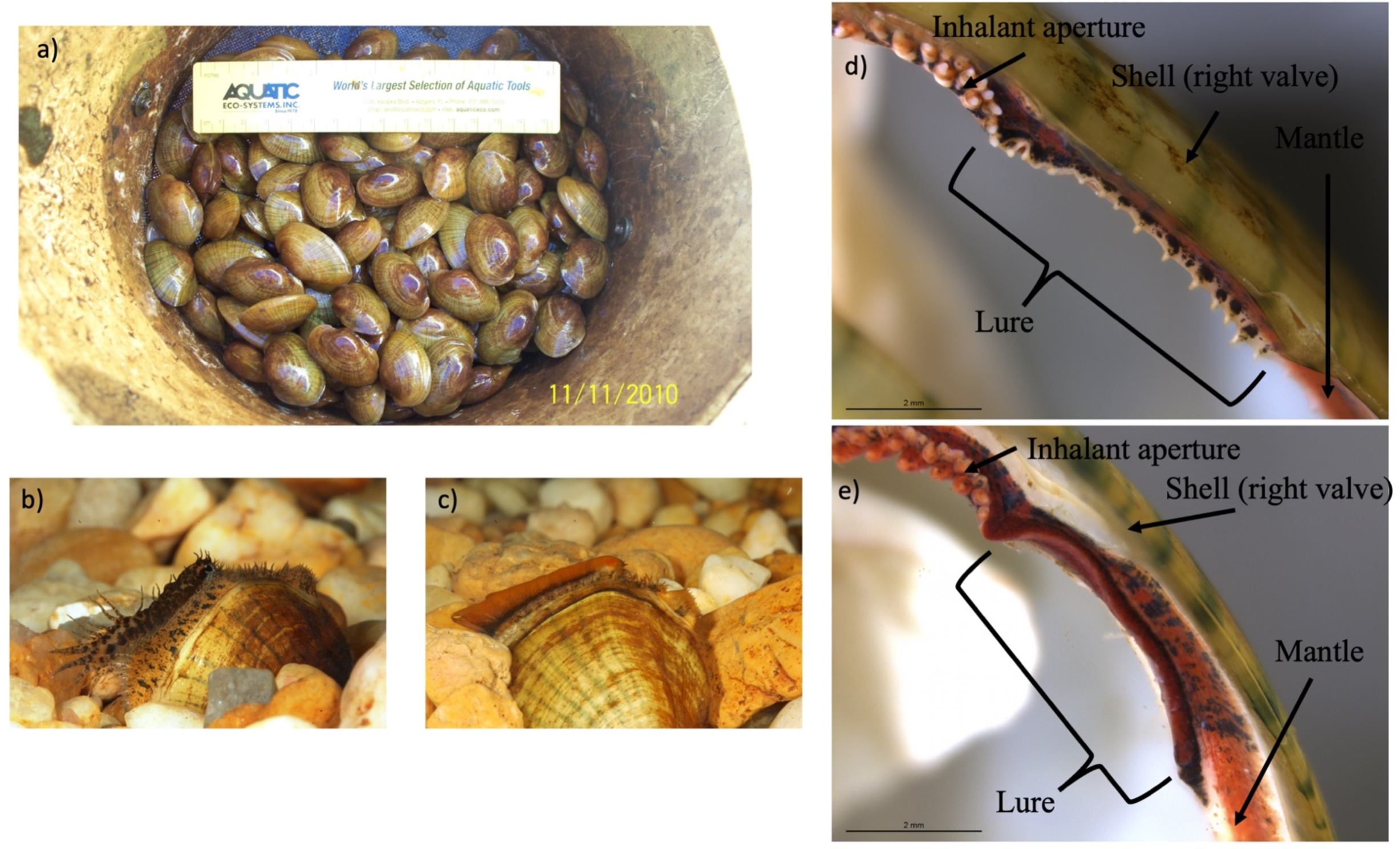
The *Lampsilis fasciola* brood raised at the Alabama Aquatic Biodiversity Center from a wild, gravid female, with a “worm-like” mantle lure (Supplementary Figure 3), sampled from the Paint Rock River (Figure 2c) in June 2009. Panel a) shows juvenile members of the brood after ∼16 months in culture. Panels b) and c) show single, sexually maturing, females after ∼2 years of culture. The young female in b) displayed a developing “darter-like” mantle lure (with “eyespots”, mottled lateral coloration, marginal extensions, and a “tail”) whereas her full-or half-sibling in c) displayed a “worm-like” mantle lure (lacking the “darter” characteristics and having orange pigmentation with a black underlay). Panels d) and e) respectively show photomicrographs, taken with a dissecting microscope, of 95% ethanol-preserved rudimentary “darter-like” and “worm-like” lures from two additional brood members, part of a 50-individual subsample preserved in 2011.

### Phylogenomic analyses

Genomic DNA was extracted from tissue samples using E.Z.N.A. Mollusk DNA kit (Omega Bio-Tek, Norcross, GA) according to manufacturer’s instructions and then stored at - 80°C. The quality and quantity of DNA extractions were assessed using a Qubit 2.0 Fluorometer (Life Technologies, Carlsbad, CA) and ddRADseq libraries were prepared following the protocols of Peterson et al. (2012). We then used 200 ng of DNA for each library prep. This involved digestion with Eco-RI-HF and MseI (New England Biolabs, Ipswich, MA) restriction enzymes, followed by isolating 294-394 bp fragments using a Pippen Prep (Sage Science, Beverly, MA) following the manufacturer’s instructions. Prepared ddRADseq libraries then were submitted to the University of Michigan’s DNA sequencing core and run in four different lanes using 150 bp paired-end sequencing on either an Illumina HiSeq 2500 or Illumina novaseq shared flow cell. Two control individuals of *Lampsilis fasciola* were run in each lane and reads for both individuals clustered together in every analysis with 100% bootstrap support, indicating no lane effects on clustering among individuals. Raw demultiplexed data were deposited at genbank under the bioproject ID PRJNA985631 with accession numbers SAMN35800743-SAMN35800847. Individuals included in phylogenomic analyses can be found in Table 1, and museum ID numbers can be found in Supplementary Table 1.

**Table 1:** Raw reads, total clusters, and total loci in assembly from the ddRAD sequencing are displayed for each genotyped sample of *Lampsilis fasciola* and of the outgroup taxa. Individual *Lampsilis fasciola* lure phenotype designation followed Zanatta et al. (2007). Museum ID numbers can be found in Supplementary Table 1.

| Sample Name | Lure Phenotype | Raw reads | Total clusters | Average clustering depth | Loci in assembly |
| --- | --- | --- | --- | --- | --- |
| L_fasciola_AL_brood_1 | Worm-like | 258664 | 97681 | 2.14 | 483 |
| L_fasciola_AL_brood_2 | Darter-like | 5201836 | 1120710 | 3.28 | 25686 |
| L_fasciola_AL_brood_3 | Worm-like | 5492519 | 1126749 | 3.4 | 25703 |
| L_fasciola_AL_brood_4 | Darter-like | 2429494 | 632254 | 2.84 | 21398 |
| L_fasciola_AL_brood_5 | Worm-like | 3152003 | 760260 | 3.02 | 23761 |
| L_fasciola_AL_brood_6 | Darter-like | 3212851 | 810898 | 2.87 | 23434 |
| L_fasciola_AL_brood_7 | Darter-like | 3649891 | 593765 | 4.22 | 25363 |
| L_fasciola_AL_brood_8 | Darter-like | 4869307 | 1462723 | 2.29 | 19089 |
| L_fasciola_AL_brood_9 | Worm-like | 3158818 | 718169 | 3.08 | 23033 |
| L_fasciola_AL_brood_10 | Darter-like | 4000321 | 915881 | 3.12 | 24916 |
| L_fasciola_AL_brood_11 | Worm-like | 5679854 | 1171842 | 3.35 | 25770 |
| L_fasciola_AL_brood_12 | Darter-like | 4212783 | 979265 | 3.04 | 24693 |
| L_fasciola_AL_brood_13 | Worm-like | 1300563 | 399134 | 2.51 | 12145 |
| L_fasciola_AL_brood_14 | Darter-like | 4100372 | 1043360 | 2.79 | 23521 |
| L_fasciola_AL_brood_15 | Darter-like | 5804293 | 1412102 | 2.91 | 25570 |
| L_fasciola_AL_brood_16 | Worm-like | 1555906 | 427061 | 2.7 | 14099 |
| L_fasciola_AL_brood_17 | Darter-like | 2073968 | 598680 | 2.59 | 13668 |
| L_fasciola_AL_brood_18 | Worm-like | 6919783 | 1574429 | 3.08 | 25811 |
| L_fasciola_AL_brood_19 | Darter-like | 3434210 | 829507 | 2.94 | 23708 |
| L_fasciola_AL_brood_20 | Darter-like | 4778853 | 994416 | 3.35 | 25500 |
| L_fasciola_AL_brood_21 | Worm-like | 2462560 | 590095 | 2.91 | 20588 |
| L_fasciola_AL_brood_22 | Worm-like | 6600876 | 1406451 | 3.26 | 26080 |
| L_fasciola_AL_brood_23 | Darter-like | 7090859 | 1628965 | 3.06 | 25932 |
| L_fasciola_AL_brood_24 | Worm-like | 4546435 | 1061394 | 3 | 24174 |
| L_fasciola_AL_brood_25 | Worm-like | 5379577 | 1135906 | 3.35 | 25703 |
| L_fasciola_AL_brood_26 | Worm-like | 5592652 | 1501130 | 2.67 | 23965 |
| L_fasciola_AL_brood_27 | Worm-like | 4893957 | 825855 | 4.09 | 25924 |
| L_fasciola_AL_brood_28 | Darter-like | 2596873 | 519103 | 3.59 | 22103 |
| L_fasciola_AL_brood_29 | Darter-like | 3401334 | 883485 | 2.87 | 21377 |
| L_fasciola_AL_brood_30 | Worm-like | 3876395 | 1014133 | 2.8 | 22072 |
| L_fasciola_AL_brood_31 | Worm-like | 5391442 | 1246528 | 3.07 | 25009 |
| L_fasciola_AL_brood_32 | Darter-like | 4365005 | 1084596 | 2.85 | 23030 |
| L_fasciola_AL_brood_33 | Darter-like | 5116507 | 1117916 | 3.16 | 24667 |
| L_fasciola_AL_brood_34 | Darter-like | 7480755 | 1601100 | 3.19 | 26163 |
| L_fasciola_AL_brood_35 | Darter-like | 8121426 | 1825135 | 3.02 | 25972 |
| L_fasciola_AL_brood_36 | Darter-like | 5521997 | 1414238 | 2.78 | 24163 |
| L_fasciola_AL_brood_37 | Darter-like | 6562641 | 1579514 | 2.88 | 25476 |
| L_fasciola_AL_brood_38 | Darter-like | 6303766 | 1596624 | 2.76 | 24448 |
| L_fasciola_AL_brood_39 | Darter-like | 6206795 | 1488925 | 2.91 | 24648 |
| L_fasciola_AL_brood_40 | Darter-like | 8630897 | 1891164 | 3.11 | 26176 |
| L_fasciola_AL_brood_41 | Darter-like | 7293683 | 1716571 | 2.95 | 25604 |
| L_fasciola_AL_brood_42 | Darter-like | 4896252 | 1193262 | 2.88 | 22829 |
| L_fasciola_AL_brood_43 | Darter-like | 6098052 | 1471714 | 2.9 | 25074 |
| L_fasciola_AL_brood_44 | Darter-like | 7495994 | 1698871 | 3.04 | 25701 |
| L_fasciola_AL_brood_45 | Darter-like | 3937758 | 670698 | 4.06 | 24947 |
| L_fasciola_AL_brood_46 | Darter-like | 6370942 | 1343655 | 3.26 | 25855 |
| L_fasciola_AL_brood_47 | Darter-like | 5542864 | 1318463 | 2.96 | 24550 |
| L_fasciola_AL_brood_48 | Darter-like | 6313913 | 1469606 | 2.98 | 24983 |
| L_fasciola_AL_brood_49 | Darter-like | 3163000 | 789239 | 2.9 | 24776 |
| L_fasciola_AL_brood_50 | Darter-like | 1728370 | 548529 | 2.35 | 17837 |
| L_fasciola_Huron_5 | Darter-like | 953302 | 259898 | 2.8 | 10996 |
| L_fasciola_Huron_6 | Worm-like | 1682931 | 362706 | 3.31 | 16809 |
| L_fasciola_Huron_7 | Worm-like | 746944 | 157212 | 3.29 | 10644 |
| L_fasciola_Huron_8 | Worm-like | 1899689 | 402515 | 3.25 | 16584 |
| L_fasciola_Huron_9 | Darter-like | 1213655 | 293090 | 2.97 | 11818 |
| L_fasciola_Huron_10 | Darter-like | 7775910 | 1275602 | 3.87 | 22035 |
| L_fasciola_Huron_11 | Darter-like | 1533281 | 295767 | 3.55 | 15386 |
| L_fasciola_NC_1 | Darter-like | 1308813 | 254002 | 3.61 | 11873 |
| L_fasciola_NC_2 | Darter-like | 4862573 | 852380 | 3.77 | 18321 |
| L_fasciola_NC_3 | Darter-like | 663874 | 165869 | 2.95 | 9960 |
| L_fasciola_NC_4 | Darter-like | 2610453 | 465228 | 3.76 | 13790 |
| L_fasciola_NC_5 | Darter-like | 6927947 | 1459334 | 3.05 | 20804 |
| L_fasciola_NC_6 | Darter-like | 1051195 | 202171 | 3.27 | 12415 |
| L_fasciola_NC_7 | Darter-like | 1948092 | 382878 | 3.61 | 17101 |
| L_fasciola_NC_8 | Darter-like | 3475751 | 669278 | 3.69 | 20683 |
| L_fasciola_NC_9 | Darter-like | 5693936 | 1634946 | 2.46 | 22325 |
| L_fasciola_NC_10 | Darter-like | 2175381 | 464794 | 3.38 | 17094 |
| L_fasciola_NC_11 | Darter-like | 2189933 | 516643 | 3.05 | 17580 |
| L_fasciola_Redo_1 | Darter-like | 1455864 | 327622 | 2.62 | 13478 |
| L_fasciola_Redo_2 | Darter-like | 1839020 | 436418 | 2.43 | 13181 |
| L_fasciola_Raisin_2 | Darter-like | 8235827 | 1716137 | 3.29 | 25555 |
| L_fasciola_Raisin_3 | Darter-like | 6032935 | 1488448 | 2.85 | 25006 |
| L_fasciola_Raisin_4 | Darter-like | 12947164 | 3587458 | 2.45 | 25245 |
| L_fasciola_Raisin_1 | Darter-like | 6639384 | 1086218 | 3.97 | 23458 |
| L_fasciola_Raisin_5 | Darter-like | 10059843 | 1997619 | 3.41 | 25363 |
| L_fasciola_Raisin_6 | Darter-like | 8019689 | 1847955 | 3.01 | 25769 |
| L_fasciola_Raisin_7 | Darter-like | 3816242 | 681697 | 3.95 | 24606 |
| L_fasciola_Raisin_8 | Darter-like | 6117037 | 1282299 | 3.27 | 22439 |
| L_fasciola_Raisin_9 | Worm-like | 5170380 | 775979 | 4.64 | 25798 |
| L_fasciola_Raisin_10 | Darter-like | 761451 | 176858 | 3.14 | 11477 |
| L_fasciola_Raisin_11 | Worm-like | 7140657 | 1670143 | 2.97 | 25519 |
| L_fasciola_Raisin_12 | Darter-like | 890521 | 203114 | 2.91 | 10582 |
| L_fasciola_Raisin_13 | Darter-like | 1071361 | 225030 | 3.47 | 13512 |
| L_fasciola_Raisin_14 | Darter-like | 3644379 | 946273 | 2.82 | 21995 |
| L_fasciola_Raisin_15 | Darter-like | 3578043 | 482446 | 5.04 | 17514 |
| L_fasciola_Raisin_16 | Darter-like | 2351544 | 114072 | 14.25 | 516 |
| L_fasciola_Raisin_17 | Darter-like | 5272816 | 1304726 | 2.87 | 23305 |
| L_fasciola_Huron_1 | Worm-like | 13366692 | 4050829 | 2.26 | 17555 |
| L_fasciola_Huron_2 | Darter-like | 2819896 | 928226 | 2.24 | 20205 |
| L_fasciola_Huron_3 | Darter-like | 662275 | 186602 | 2.66 | 7653 |
| L_fasciola_Huron_4 | Darter-like | 4792093 | 855457 | 3.88 | 24512 |
| L_fasciola_AL_mom_1 | Darter-like | 8095030 | 1840917 | 2.95 | 25420 |
| L_fasciola_AL_mom_2 | Darter-like | 10329331 | 3504027 | 2.03 | 24488 |
| L_fasciola_AL_mom_3 | Darter-like | 10384477 | 2987559 | 2.34 | 25056 |
| L_fasciola_Huron_12 | Worm-like | 6906349 | 1672394 | 2.87 | 25281 |
| L_fasciola_Huron_13 | Worm-like | 6955496 | 1670627 | 2.88 | 25593 |
| L_fasciola_Raisin_18 | Worm-like | 5506215 | 1301878 | 3 | 25373 |
| L_fasciola_Raisin_19 | Worm-like | 6611596 | 1524682 | 3.03 | 25604 |
| L_fasciola_Raisin_20 | Worm-like | 4894495 | 1276608 | 2.74 | 24931 |
| L_fasciola_Raisin_21 | Worm-like | 8396562 | 1736736 | 3.26 | 25490 |
| L_cardium_1 |  | 6864226 | 1710220 | 2.8 | 14625 |
| L_cardium_2 |  | 4898330 | 1091622 | 3.11 | 13433 |
| L_cardium_3 |  | 7109883 | 2005565 | 2.5 | 14563 |
| L_cardium_4 |  | 4637077 | 997208 | 3.27 | 13860 |
| S_nasuta_1 |  | 4544989 | 1169260 | 2.55 | 10441 |

The alignment-clustering algorithm in ipyrad v.0.7.17 (Eaton, 2014; Eaton & Overcast, 2020) was used to identify homologous ddRADseq tags. Ipyrad is capable of detecting insertions and deletions among homologous loci, which increases the number of loci recovered at deeper evolutionary scales compared to alternative methods of genomic clustering (Eaton, 2014). Demultiplexing was performed by sorting sequences by barcode, allowing for zero barcode mismatches (parameter 15 setting 0) and a maximum of five low-quality bases (parameter 9). Restriction sites, barcodes, and Illumina adapters were trimmed from the raw sequence reads (parameter 16 setting 2), and bases with low-quality scores (Phred-score <20, parameter 10 setting 33) were replaced with an N designation. Sequences were discarded if they contained more than 5 N’s (parameter 19). Reads were clustered and aligned within each sample at an 85% similarity threshold, and clusters with a depth < 6 were discarded (parameters 11 and 12). We also varied the number of individuals required to share a locus from ∼50% to ∼75%.

We analyzed the two concatenated ddRAD-seq alignment files (50% and 75% minimum samples per locus) using maximum likelihood in RAxML v8.2.8 (Stamatakis, 2014). A general time-reversible model (Lanave et al., 1984) was used for these analyses that included invariable sites and assumed a gamma distribution. Support was determined for each node using 100 fast parametric bootstrap replications. Lure phenotype information was recorded and mapped on to the phylogenetic tree. Phylogenetic signal of lure phenotype was tested using Pagel’s λ (Pagel, 1999) in R (R Core Team, 2018) with the ‘phylobase’ package (R Hackathon et al., 2013).

### River Raisin Mantle Lure Phenotype Ratios Over Time

Mid-20^th^ century *Lampsilis fasciola* specimens collected at the Sharon Mills County Park site (Raisin River, MI; Fig. 2a) are preserved as part of the University of Michigan’s Museum of Zoology wet mollusk collection. They stem from 8 different collecting events between 1954 and 1962 (Table 2) and their presence afforded an opportunity to assess the stability of the *L. fasciola* “darter/worm” mantle lure polymorphism in that population over a six-decade time interval. All of the museum specimens, males as well as females, were examined to determine whether their fully-formed (female) or rudimentary(male) mantle lures were “darter-like” or “worm-like”. For females, this could be achieved by simple visual examination, but male lure classification required a dissecting microscope. The ratio of mantle lure phenotypes observed in the Sharon Mills County Park population was compared among mid-20th century (UMMZ preserved females and males) and 2017 (field photographs and videos of displaying females) samples using a Fisher’s exact test, implemented in R.

**Table 2:** Summary of the sex and lure phenotypes of all 57 University of Michigan Museum of Zoology *Lampsilis fasciola* individuals present in 8 separate mid-20^th^ century collections made from the River Raisin at Sharon Mills County Park (Fig. 2a).

| Collection Date | Male darter-like | Male worm-like | Female darter-like | Female worm-like | Total darter-like | Total worm-like | Total |
| --- | --- | --- | --- | --- | --- | --- | --- |
| 9/30/54 | 2 | 1 | 5 | 1 | 7 | 2 | 9 |
| 5/21/58 | 0 | 0 | 4 | 0 | 4 | 0 | 4 |
| 7/28/59 | 0 | 0 | 2 | 0 | 2 | 0 | 2 |
| 4/24/62 | 3 | 0 | 3 | 0 | 6 | 0 | 6 |
| 5/10/62 | 5 | 1 | 4 | 1 | 9 | 2 | 11 |
| 6/19/62 | 4 | 2 | 1 | 0 | 5 | 2 | 7 |
| 6/25/62 | 3 | 1 | 2 | 0 | 5 | 1 | 6 |
| 7/20/62 | 5 | 1 | 5 | 1 | 10 | 2 | 12 |

### Putative Lure Mimicry Models

Population-specific putative model species for the *Lampsilis fasciola* mantle lure mimicry system were investigated at the River Raisin Sharon Mills County Park study site (Figure 2) in part because of the availability of a comprehensive ecological survey of Raisin River fishes (Smith, Taylor & Grimshaw, 1981). “Darters” – members of the speciose North American subfamily Etheosomatidae – have been implictly identified as models for the predominant “darter-like” mantle lure phenotype (Zanatta, Fraley & Murphy, 2007), and they are preyed upon by *Micropterus dolomieu* (Surber, 1941; Robertson & Winemiller, 2001; Murphy et al., 2005), *L. fasciola’s* primary fish host (Zale & Neves, 1982; McNichols, 2007; Morris et al., 2009; McNichols, Mackie & Ackerman, 2011; VanTassel et al., 2021). Ten species of Etheosomatidae occur in the River Raisin, as does *M. dolomieu* (Smith, Taylor & Grimshaw, 1981).

River Raisin gravid female *Lampsilis fasciola* engage in mantle lure displays from May-August. During the summer of 2017, a total of 27 different displaying females were photographed along a 150m stretch downstream of the dam at Sharon Mills county park using an Olympus Tough TG-6 underwater camera. The lures were first categorized into broad groupings based on visual similarity (in terms of morphology and coloration; Supplementary Figure 2) and these groupings were then used to identify putative host prey fish model species from those present in the River Raisin drainage (Smith, Taylor & Grimshaw, 1981), in terms of convergent size, shape, and coloration. Putative model species were further assessed based on their relative local abundance (Smith *et al*., 1981) and on their range overlap with both mimic and receiver. Geographic ranges of *L. fasciola,* the primary host *M. dolomieu*, and each prospective model species were carefully produced by hand in Arcgis software (ESRI, 2022), and the overlap between *L. fasciola, M. dolomieu,* and each putative model species was assessed using Arcgis software.

### Behavioral Analyses

Standardized video recordings of 30 mantle lure-displaying female *L. fasciola* were recorded using a Go Pro Hero 6 camera in the Summer of 2018 at the two different southeastern Michigan study sites: Sharon Mills County Park (River Raisin) and Hudson Mills Metropark (Huron River). An additional 4 video recordings of the lure behavior for the sympatric congener *Lampsilis cardium* were collected from the Sharon Mills site to assess interspecific variability in lure behavior. Recordings were captured from a top-down perspective during daylight hours using a standardized frame that included a metric ruler and a Casio TX watch to record date, time, and water temperature data within the video frame. For each displaying female, videos of the lure movements were recorded for 10 minutes at 120 frames-per-second. Analysis of the videos involved manually recording mantle lure movements for 20,000 frames, starting at 5,000 frames to to avoid any initial setup effects on mussel display behavior. The frame numbers when an individual movement began and ended were noted, and movements of the left and the right mantle lure flaps were assessed seperately.

To quantitatively assess behavioral differences among samples, gait analysis diagrams were created in R for each displaying mussel. Averages and standard deviations for the time intervals between lure undulations were calculated for each individual, as well as speed of undulation and proportion of movements synchronized. A Kruskal-Wallis test was used to test for overall differences among lure groups (*L. fasciola* “darter-like”, *L. fasciola “*worm-like”, and *L. cardium*), and pairwise Wilcoxon Signed rank tests were used to compare groups directly with a Bonferroni *P* value adjustment to correct for multiple tests. A Spearman correlation was used to test for water temperature on time interval between lure undulations.

To further explore differences in lure behavior among groups, we used a General Linear Mixed Model (GLMM), with sample ID as a random factor, to test for differences in lure movement intervals. The GLMM appoach, unlike simple mean comparisons, allows the inclusion of all lure movements for all individuals in the model. Because displaying mussels all varied in the number of lure movements recorded over 20,000 frames analyzed, a dataset of 1000 random bootstrap values was constructed for each individual by randomly sampling values, with replacement. Models were fitted using the ‘lmerTest’ package in R, and Satterthwaite’s Method (Satterthwaite, 1946; Kuznetsova, Brockhoff & Christensen, 2017) was used to test for significance of fixed effects of lure phenotype on the interval between lure undulations.

## RESULTS

### Captive Brood

Mantle lure microphotographs (Figure 3d, e) of all 50 available AABC-cultured and 95% ethanol-preserved *Lampsilis fasciola* of the same maternal brood are individually presented in Supplementary Figure 4. These specimens were 2 years post-metamorphosis when preserved in 2011 and independent classification of their mantle lure phenotypes concurred the ratio of “darter-like” to “worm-like” phenotypes was respectively 33/17 for this brood subsample.

### ddRAD-seq and Phylogenomic analyses

Genomic sequencing returned raw reads ranging from 258,664 to 13,366,692 per individual across the 108 unionid specimens included in the analyses comprising samples of the ingroup *Lampsilis fasciola*, sourced from 4 different populations, along with outgroups *Lampsilis cardium* and *Sagittunio nasuta*. Mean coverage depth for the 85% clustering threshold ranged from 2.03 (L_fasciola_AL_mom_2) to 14.25 (L_fasciola_Raisin_16; Table 1). Between 28,725 and 16,161 homologous loci were identified across the two best ddrad datasets (85%-50% and 85%-75% respectively) and the number of loci recovered was generally consistent among all samples.

The maximum likelihood tree produced by RAxML (Supplementary Figure 5) recovered the following ingroup*/*outgroup topology: (*S. nasuta* (*L. cardium, L. fasciola*)) with outgroup branch lengths greatly exceeding those of the ingroup. To optimize the legibility of ingroup relationships, a compressed, color-coded graphic excluding *S. nasuta* was constructed (Figure 4). A nested series of phylogenetic relationships was recovered for the four *L. fasciola* fluvial populations with the two Michigan drainages being paraphyletic: (Little Tennessee River (Paint Rock River (River Raisin (River Raisin, Huron River))). The ingroup topology also showed evidence of within-population genealogical relationships with all Paint Rock River brood members forming an exclusive clade (Figure 4).

**Figure 4:**
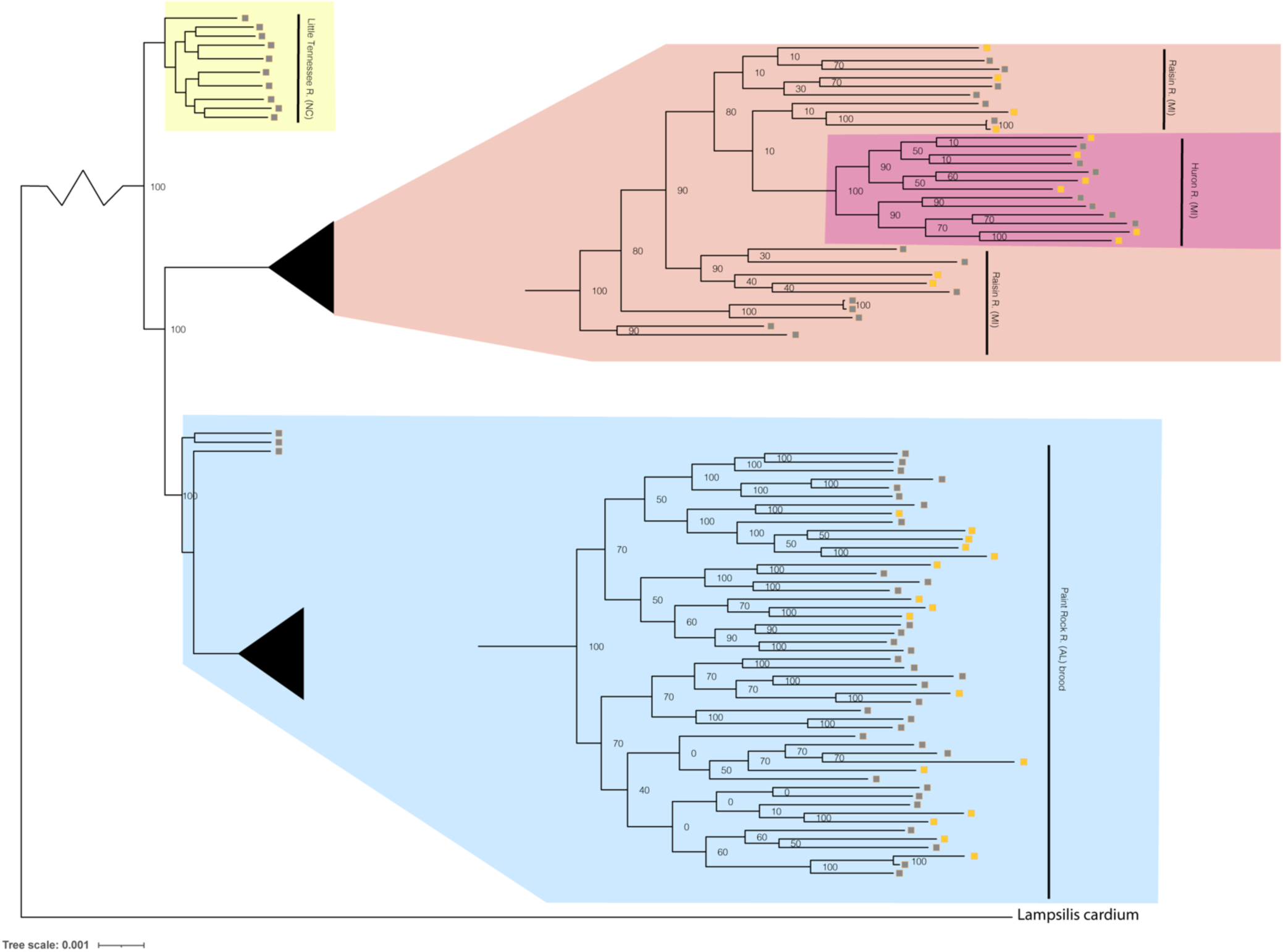
Phylogenomic tree of 96 *Lampsilis fasciola* individuals created in RAxML using 28,735 concatenated ddRAD-seq loci. Gravid, lure-displaying females sampled from two Michigan drainages, River Raisin (Fig. 2a) and Huron River (Fig. 2b), are respectively highlighted in peach and pink. Specimens sampled from the Paint Rock River, Alabama (Fig. 2c) are highlighted in blue and consisted of three gravid, lure-displaying females, in addition to 50 larval brood members raised at the Alabama Aquatic Biodiversity Center in the zoomed-in tip clade. Gravid, lure-displaying females sampled from the Little Tennessee River in North Carolina (Fig. 2d) are highlighted in yellow. The respective primary mantle lure phenotypes – “darter-like” or “worm-like” – of all *L. fasciola* individuals is indicated by gray and yellow boxes, respectively.

The respective primary mantle lure phenotypes – “darter-like” or “worm-like” – of all 92 *Lampsilis fasciola* ingroup individuals is indicated in Figure 4. Note that 3 of the 4 population samples – Little Tennessee River, River Raisin and Huron River – were exclusively composed of mantle-lure displaying wild females, and the latter two samples were polymorphic in mantle lure composition. Regarding the Paint Rock River sample, polymorphic lures were restricted to the 50 captive-raised AABC brood members sourced from a gravid, wild female in 2009 (not included in the analyses). The ingroup phylogeny (Figure 4) contained two polymorphic mantle lure clades, one composed of both Michigan populations (River Raisin and Huron River), the other consisting only of the AABC brood, and both clades had individuals of either lure phenotype interspersed across their respective topologies. Little phylogenetic signal associated with either primary mantle lure phenotype (11 = 0.21; *P* = 0.13).

### Phenotypic Ratios Over Time

Table 2 summarizes the sex and primary lure phenotypes of 57 *Lampsilis fasciola* specimens collected from 1954-1962 at the River Raisin Sharon Mills County Park study site (Figure 2a) and preserved in the University of Michigan Museum of Zoology’s wet mollusk collection (Figure 5b, c). These historical samples had a collective “darter-like” to “worm-like” ratio of 48:9, with 15.8% of individuals having the less common “worm-like” mantle lure phenotype. Figure 5a contrasts the mid-20^th^ century lure phenotype ratios with a contemporary (2017) estimate in that same population, based on photographic recordings of 27 displaying females. The contemporary estimate was 23:4, with 14.8% of individuals having the less common “worm-like” mantle lure phenotype. A Fisher’s exact test found no significant differences between these two temporal estimates of relative mantle lure phenotype frequencies in the River Raisin Sharon Mills County Park *L. fasciola* population (*P* = 1).

**Figure 5:**
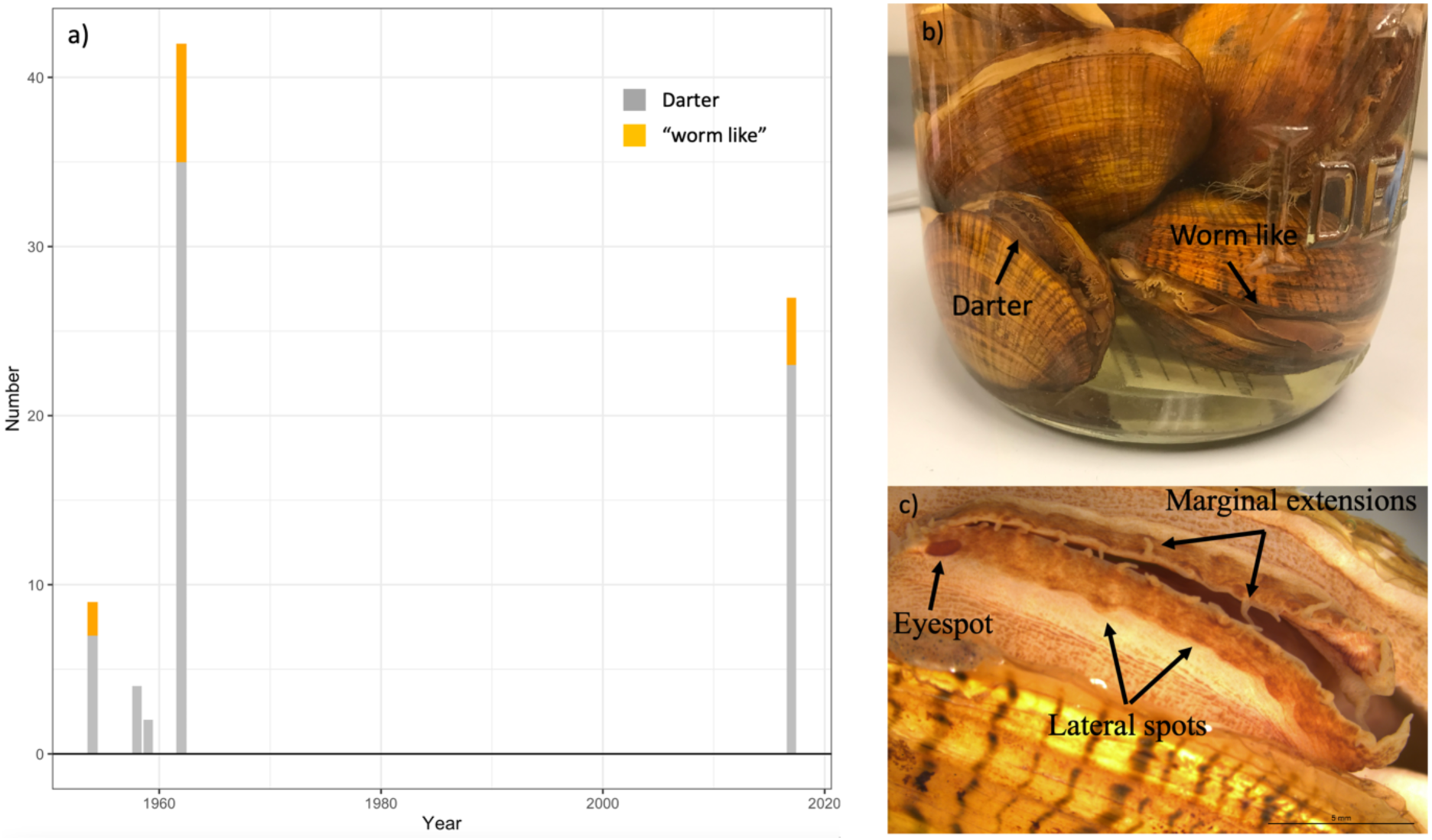
a) The observed frequency of River Raisin *Lampsilis fasciola* primary mantle lure phenotypes (“darter-like” vs “worm-like”) at the Sharon Mills County Park study site during two different time periods. The 1954-1962 data were obtained from the University of Michigan Museum of Zoology (UMMZ) collection specimens, both female and male. The 2017 data were based on field observations of displaying females. b) a jar of preserved UMMZ Sharon Mills specimens showing a “darter-like” and a “worm-like” mantle lure. c) a “eyespot”, lateral pigmented blotches, and marginal extensions.

### Putative Raisin River Lure Mimicry Models

The field photographs of 27 displaying female *L. fasciola* mantle lures in the Raisin River Sharon Mills County Park population in 2017 (Supplementary Figure 2) were categorized into either “darter-like” (Zanatta, Fraley & Murphy, 2007) or “worm-like” (McNichols, 2007), as summarized in the Materials & Methods section. In addition to the specific features that separate these two primary mantle lure phenotypes (presence/absence of “eyespots” mottled pigmentation, marginal extensions and a “tail”), “darter-like” lures exhibited a much higher degree of variation than did “worm-like” lures, both within populations and across the species range. The latter lure phenotype exhibited a relatively simple, uniform morphology combined with a bright orange coloration underlain with a black basal stripe phenotype in Michigan (Figure 6f-h), in Alabama (Fig. 6i, j), and in North Carolina populations (Fig. 2A in Zanatta et al, 2007). In contrast, Raisin River “darter-like” mantle lures exhibited individual-level variation that was sometime quite marked, especially in details of their pigmentation, and to a more limited degree in their marginal extensions (Figures 6a-d; Supplementary Figure 2). Among individual variation was most pronounced for inter-population camparisons, e.g., see the much larger “tail” in the lure displaying Paint Rock River, Alabama specimen shown in Figure 6e, and also the wider range of phenotypes present in North Carolina populations (Fig. 2b-d in Zanatta et al, 2007).

**Figure 6:**
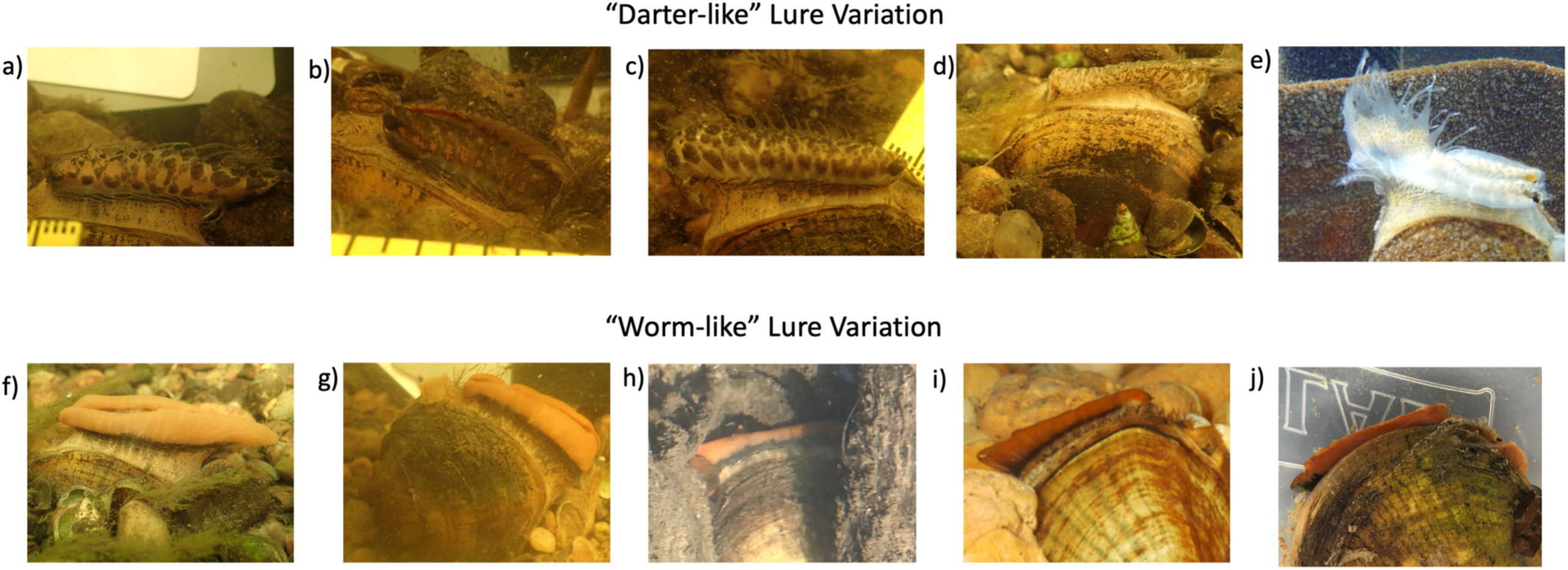
This panel displays some of the variability in lure phenotype, both within a population and across the range of *Lampsilis fasciola*. a-d are “darter-like” Raisin River (MI) lures photographed in the field at Sharon Mills County Park. e) depicts a “darter-like” lure displayed by a Paint Rock River (AL) female. f-h show field photographs of “worm-like” lures displayed by three Sharon Mills females, with specimen h being a younger adult. i,j are photographs of two captive AABC specimens, with “worm-like” lures, sourced from the Paint Rock River. The former photo (i), taken in 2011, shows a young (2-year old) female, a member of the captive brood, displaying her lure, and the latter photo (j) is of a female field-sampled in 2022, and showing a partially retracted mantle lure.

Despite the considerable individual variation among the 24 photographed Raisin River “darter-like” mantle lures (Supplementary Figure 2), it was possible to identify some shared phenotypic motifs, especially in pigmentation pattern, and to informally categorize 23/24 mantle lures with those shared motifs into 4 general groupings. Group 1 “darter-like” mantle lures were characterized by prominent, chevron-like, darker pigmented blotches, spaced regularily along the flanks of the lure, over a lighter background coloration (Figure 6a). This general pattern occurred in 7/24 Raisin River “darter-like” lures examined. Group 2 was rarer (3/24 individuals) and consisted of a darker background coloration with large orange blotches spaced regularily along the lure flanks, some divided into “dorsal” and “ventral” elements (Figure 6b). Group 3 (9/24 individuals) lures were characterized by prominent dark lateral maculation spatially divided into a “ventral” pattern of larger, regularly spaced blotches and a “dorsal” pattern of more numerous, irregular blotches of different sizes (Figure 6c). Finally, Group 4 (3/24 individuals) lures were characterized by an evenly-dispersed, fine grained freckling of numerous pigmented spots over a lighter background (Figure 6d).

To explore putative model species for the four *Lampsilis fasciola* Raisin River “darter-like” mantle lure groupings (Figure 6a-d), potential matches (in terms of size, shape and coloration) were sought among the 10 species of Etheosomatidae that occur in the River Raisin (Smith *et al*., 1981), many of which display pronounced sexual dimorphism in body coloration (Kuehne and Barbour, 2014). The best apparent matches, depicted in Figure 7, are as follows: Group 1 (Fig. 6a)-*Etheostoma blennioides* (female coloration), Group 2 (Fig. 6b)-*Etheostoma exile* (male coloration), Group 3 (Fig. 6c)-*Percina maculata* (male and female coloration) and Group 4 (Fig. 6d)-*Etheostoma microperca* (female coloration).

**Figure 7:**
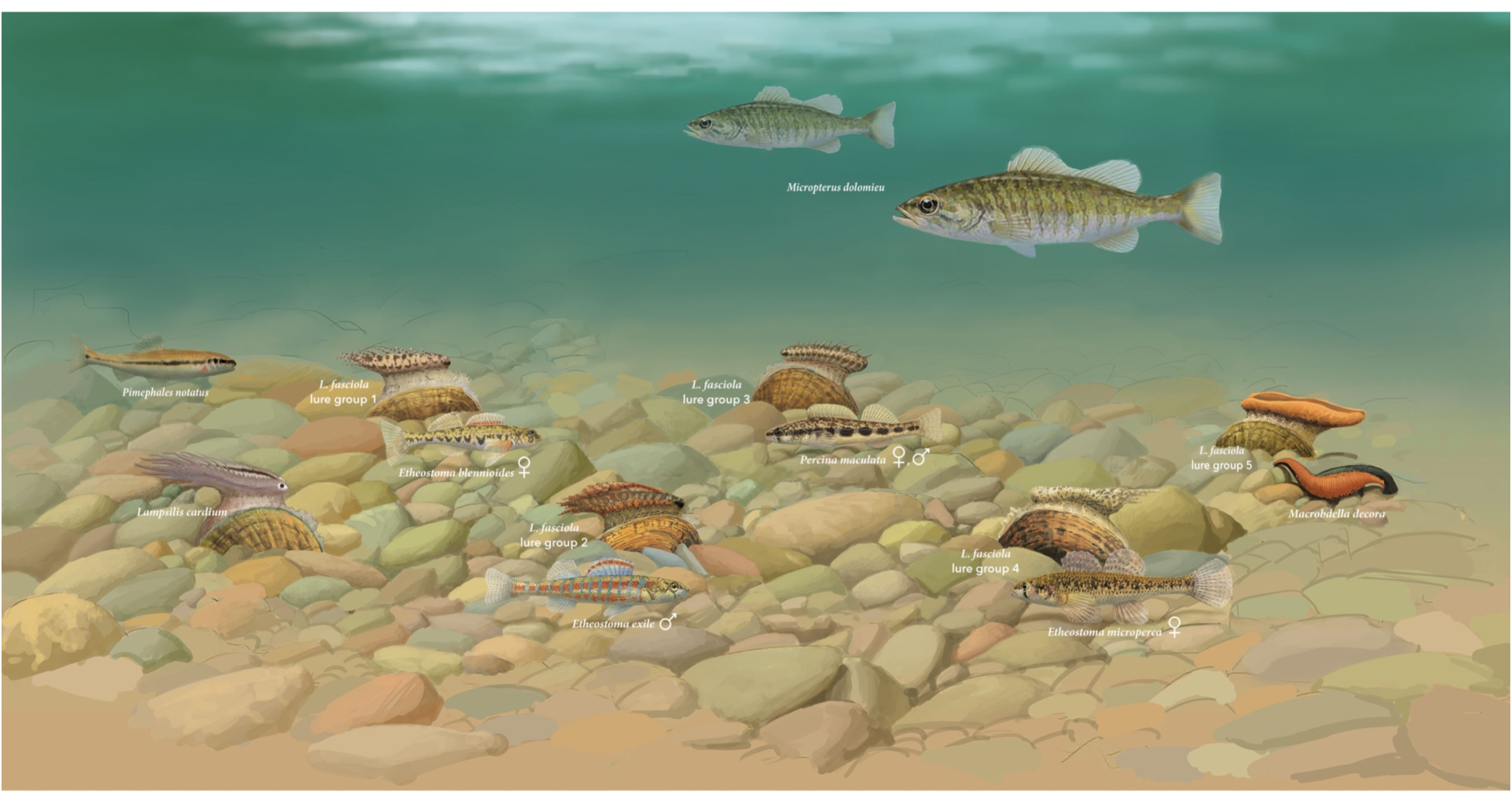
A hypothetical Raisin River (Michigan) benthic assemblage showing displaying exemplars of the putative 5 main *Lampsilis fasciola* mimetic mantle lure groups (Figure 6a-d, f) present at the Sharon Mills County Park study site, together with their respective model species, and their primary receiver/fish host, *Micropterus dolomieu*. Also shown is a displaying *Lampsilis cardium* with a “small minnow” mimetic mantle lure (Patterson et al., 2018) and its putative model, *Pimephales notatus*, the most common fish species in the River Raisin (Smith, Taylor & Grimshaw, 1981).

The distinctive color combination of the *Lampsilis fasciola* “worm-like” lure - solid orange with a black underlay (Figure 6 f-j) - did not match that of any Raisin River darter, or other Raisin River fishes (Smith, Taylor & Grimshaw, 1981). It does, however, match the coloration and size/shape, of the common North American leech, *Macrobdella decora*, which is widespread and abundant in eastern North America watersheds and typically feeds on aquatic vertebrates (Klemm, 1982; Munro et al., 1992). *M. dolomieu*, *L. fasciola*’s primary host fish, is a generalist predator with a diet of aquatic invertebrates, including leeches, in addition to small fishes (Clady, 1974) and recreational fishers frequently use live and/or artifical leeches as bait to catch this species (Cooke et al., 2022). Based on the available data, it seems that *Macrobdella decora* may be the best model species candidate for the “worm-like” (McNichols, 2007) *L. fasciola* mantle lure phenotype, and will hereafter be referred to as the leech phenotype.

The geographic range of the mimic, *Lampsilis fasciola*, is a subset of that of its receiver/host *M. dolomieu* (Figure 2), and the extent of range overlap with all 5 putative River Raisin mantle lure models were calculated using Arcgis (Table. 3) and are shown in Figure 8. Three of the five putative models -*Etheostoma blenniodes, Percina maculata* and *Macrobdella decorata* have extensive overlap with *L. fasciola*’s range, but *E. exile* and *E. microperca* are restricted to northern portions.

**Figure 8:**
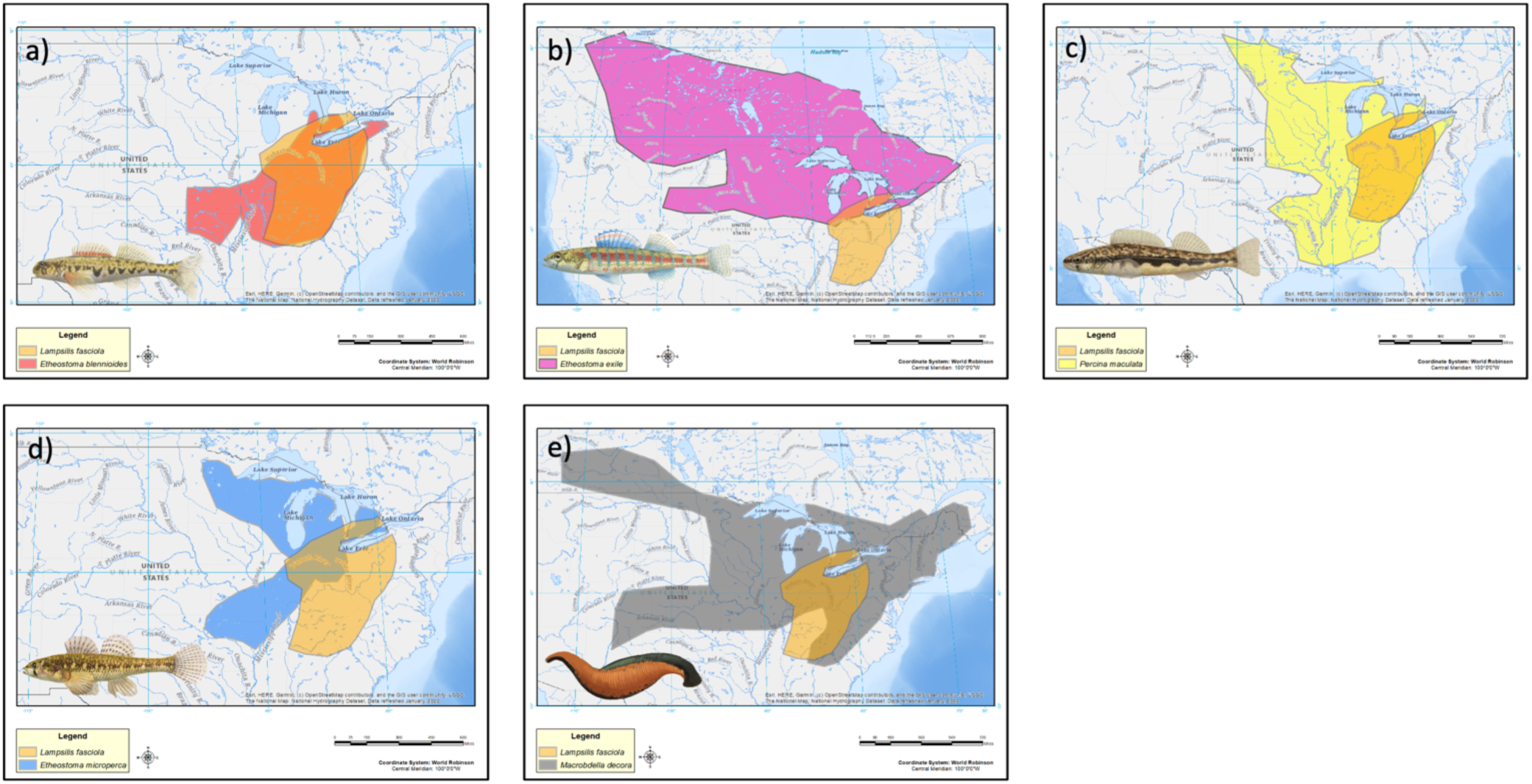
Estimated range maps for 5 proposed models for *Lampsilis fasciola* lures. Range maps for each proposed model is compared to the estimated geographic range of *Lampsilis fasciola*. a) *Etheostoma blennoides*, b) *Etheostoma exile*, c) *Percina maculata*, d) *Etheostoma microperca*, and e) *Macrobdella decora.* Model Illustrations by John Megahan.

**Table 3:** The 5 broad categories of lure phenotypes (Groups a-e) observed at the River Raisin Sharon Mills County Park population of *Lampsilis fasciola* (Fig. 2a), as well as the estimated geographic range overlap between *Lampsilis fasciola* and its 5 Raisin River putative model species.

| Type | Proposed Model | Range overlap (km<sup>2</sup>) |
| --- | --- | --- |
| Group a | *Etheostoma blennioides* | 480,731 |
| Group b | *Etheostoma exile* | 87,796 |
| Group c | *Percina maculata* | 525,772 |
| Group d | *Etheostoma microperca* | 164,539 |
| Group e | *Macrobella decora* | 419,259 |

### Behavioral Analyses

Lure movement data was extracted from video recordings of 30 *L. fasciola* and 4 *L. cardium.* Qualitatively, *L. fasciola* and *L. cardium* have very different mantle lure display behaviors. Gait diagrams show a clear distinction between both primary *L. fasciola* lure phenotypes (“darter” and “leech”) and *L. cardium*. A representative individual for each group is shown in Figure 9, and gait diagrams for all individuals can be found in Supplementary Figure 6. *L. cardium* consistently exhibited a synchronized lure undulation of both mantle lure flaps, whereas *L. fasciola* samples frequently moved left and right mantle flaps independently. Gait diagrams also qualitatively showed that while *L. fasciola* behavior is charcterized by a high level of variability in undulation interval, *L. cardium* is comparitavely much more regular in undulation interval with a steady beat frequency (Figure 9a-c).

**Figure 9:**
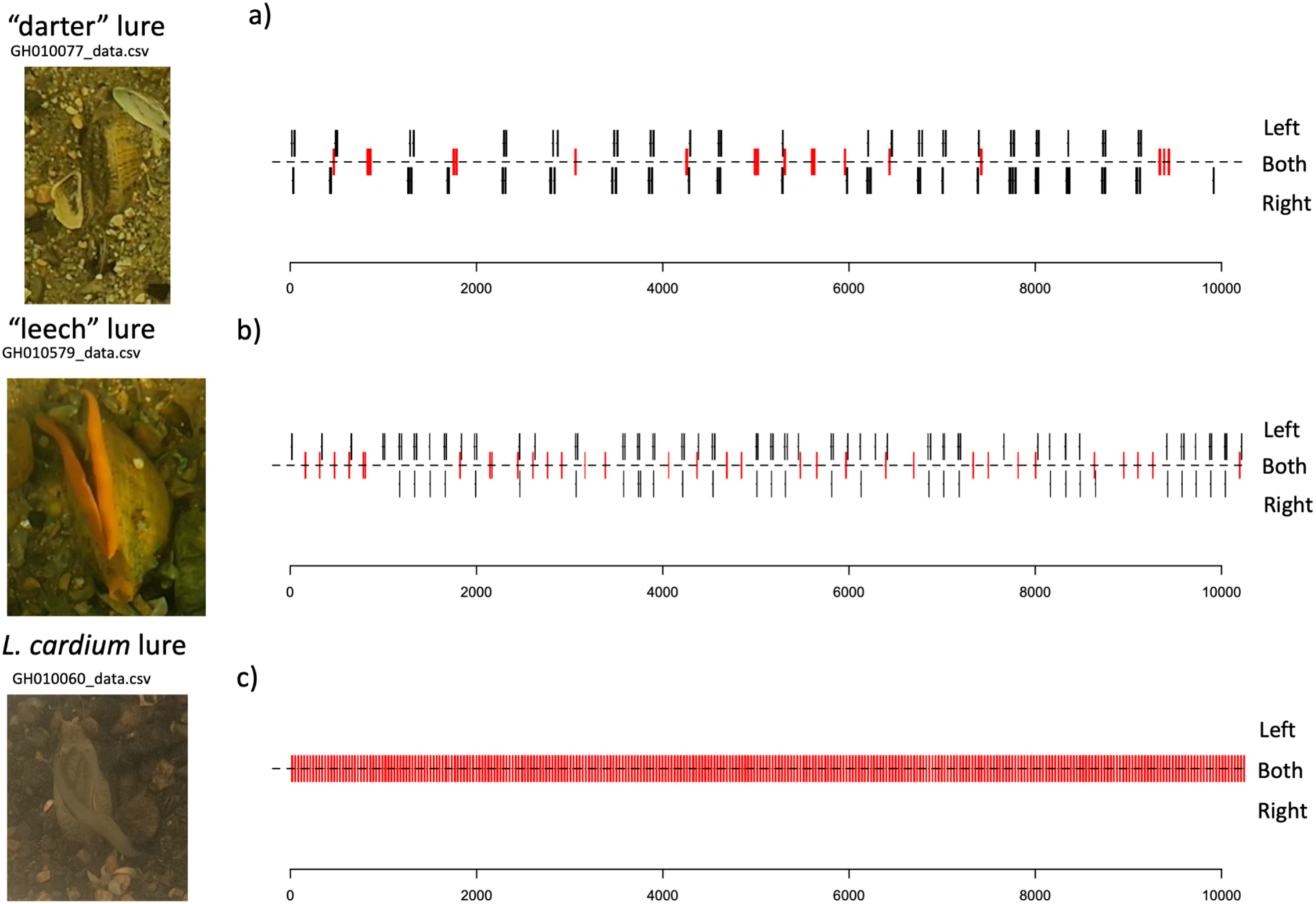
Mantle lure gait diagrams for three representative individuals that were field recorded using a Go Pro Hero 6 in 2018. Panel a) shows a *Lampsilis fasciola* “darter” lure sample, Panel b) displays a *Lampsilis fasciola* “leech” lure sample, and Panel c) shows a *Lampsilis cardium* sample. Red center lines indicate synchronized lure movement for both left and right mantle flaps, and black lines above and below the center line indicate independent left and right movements respectively. The x-axis denotes frame number (120fps).

Supplementary Table 2 details the time, date, location, temperature and summary statistics of all 34 lure display field recordings. Movements of the left and right mantle flaps in *L. fasciola* were recorded seperately. Boxplots for means and standard deviations of intervals between left lure movements are displayed in Figure 10a,b. Boxplots of mean duration of lure movements (L) are shown in Figure 10c and the proportion of synchronized lure movements are displayed in Figure 10d. Only left mantle flap movements are displayed in Figure 10a-c, however the distributions for the right mantle flap movements are highly similar. Kruskal-Wallis tests suggest highly significant differences among all groups for average inter-undulation interval and for standard deviation of inter-undulation interval (*P <* 0.001). Pairwise Wilcoxon tests revealed highly significant differences in lure inter-movement interval (left) between *L. cardium* and both *L. fasciola* groups (*P* = 0.001 and 0.001 for “darter” and “leech” respectively) and between both primary *L. fasciola* mantle lure phenotypes as well (*P* = 0.01). Pairwise Wilcoxon tests also revealed significant differences in the standard deviation of left inter-movement interval between *L. cardium* and both *L. fasciola* groups (*P* = 0.001 and 0.01 for “darter” and “leech” respectively) but not between the two *L. fasciola* mantle lure groups (*P* = 0.09). These results suggest that, in addition to the qualitative differences between *L. cardium* and *L. fasciola* lure behavior, the *L. cardium* lures are beating at a faster rate and with greater regularity. There also appears to be a slight difference in the rate of lure beats between the two *L. fasciola* mantle lure morphs, with the “darter” lure beating slightly less frequently, but with no differences in variability. No significant relationship was observed between lure inter-movement interval (left) and water temperature (Supplementary Figure 7; *P* = 0.428).

**Figure 10:**
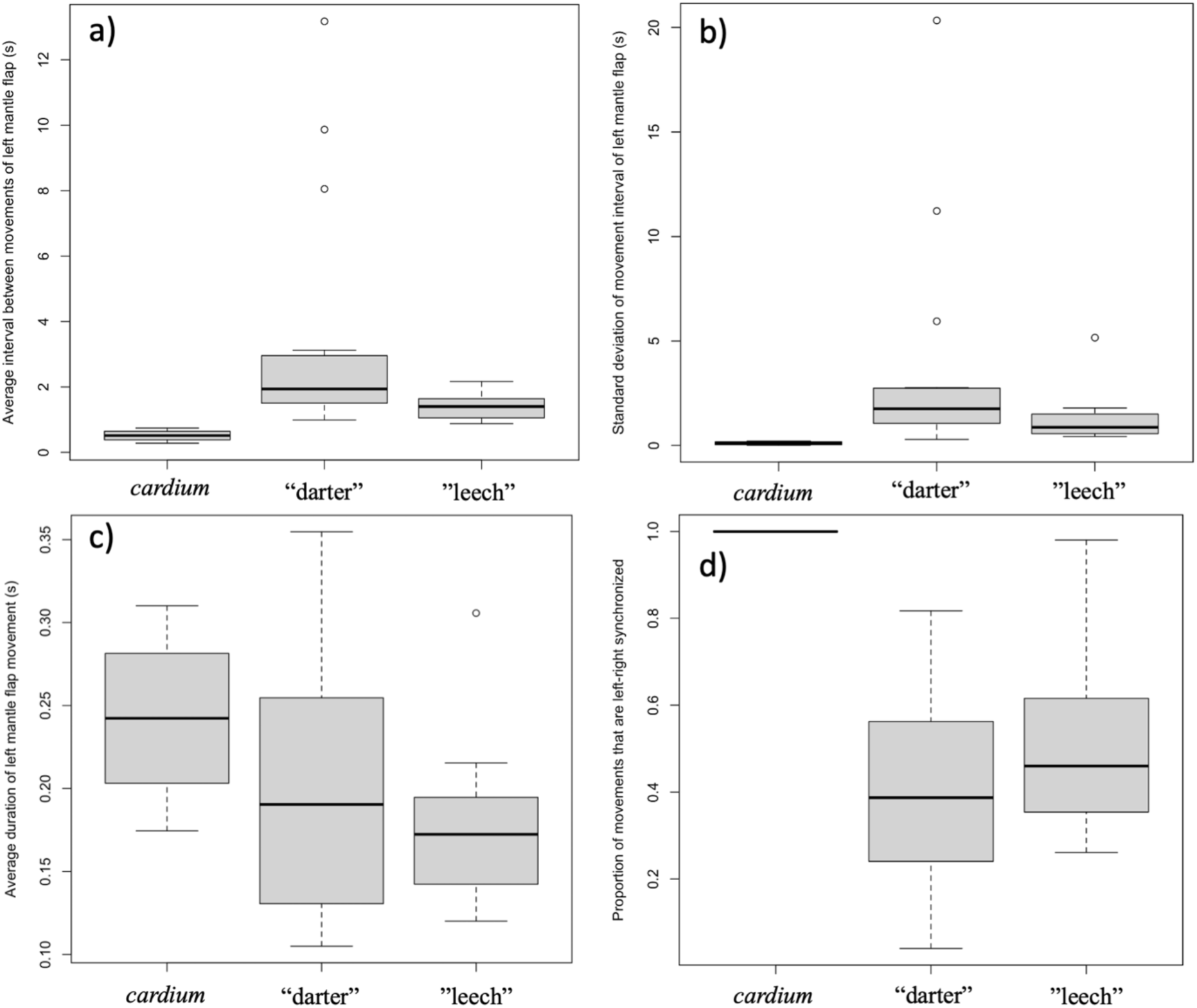
Summary boxplots from behavioral analyses of the two primary *Lampsilis fasciola* mantle lure phenotypes (“darter” vs. “leech”) and of *Lampsilis cardium*. Panel a) comparison of the mean interval between lure movements. Panel b) the mean standard deviation in lure movement interval as a proxy of variability. Panel c) the mean duration of lure movement. Panel d) the proportion of movements that are left-right synchronized. Panels a-c show means for left mantle flap movements only.

General Linear Mixed Models (GLMM) were used as an alternative analytical approach that included a large, bootstrapped dataset of lure movements. GLMM results were similar to those of the mean comparisons, with *L. cardium* individuals having faster movement frequency than either *L. fasciola* lure morphs (an estimated 0.21 seconds for *L. cardium* versus 3.2 and 1.0 seconds respectively for L. fasciola “darter” and “leech” lures). However, these fixed effects are not significant.

## DISCUSSION

Two new lines of data, phylogenomic and genetic, corroborated Zanatta et al’s. (2007) preliminary finding that the primary mantle lure morphs in *Lampsilis fasciola* (Figure 1b, c) represent a within-population polymorphism rather than cryptic taxa. In phylogenomic analyses, all three polymorphic population samples (Huron, Raisin, and Paint Rock Rivers), collectively spanning the species range (Figure 2a-c), produced tip clades that were comprehensively polyphyletic regarding lure morph type (Figure 4) and the “darter vs leech” dichotomy yielded a low estimate of phylogenetic signal (λ = 0.21). However, the phylogenomic data did reveal clear evidence of geographic structuring (Figure 4), even among regional populations with a continuous freshwater connection. For example, the Huron and Raisin drainages empty in Western Lake Erie and the Little Tennesse and Paint Rock drainages empty into the Tennessee River (see also VanTassel et al. (2021)). The Paint Rock River (AL) population was sister to the Michigan populations (Figure 4), a result consistent with phylogeographic associations of multiple other North American species, including unionid mussels and *Micropterus dolomieu*, attributed to hypothesized glacial refugia in the southern Appalachian mountains (Soltis et al., 2006; Borden & Krebs, 2009; Zanatta & Harris, 2013; Hewitt et al., 2018).

Discovery of within-brood mantle lure heterogeneity (Figure 3), apparently the first such record for unionids, confirms that the *Lampsilis fasciola* “darter-like” and “leech-like” mantle lures are true polymorphisms and provides initial, although limited, genetic insights into lure phenotype inheritance. Of the 50 available offspring, the maternal “leech” phenotype was inherited by 17, the remaining 33 had the “darter” phenotype, but none exhibited a recombinant phenotype, *e.g.*, “leech” coloration with “darter” marginal extensions or “darter” coloration without marginal extensions. Evidence of discrete, within-brood segregation of the mantle lure polymorphism implies potential control by a single genetic locus and expression of the maternal phenotype in ∼1/3rd of the offspring is inconsistent with a dominant hypothetical “leech” allele. Additional pedigree insights are currently inhibited by not knowing the number of sires that contributed to the brood: the dam was a wild-mated Paint Rock River individual. Freshwater mussel broods frequently have multiple paternity (Ferguson et al., 2013; Wacker et al., 2018) and this may well be the case also for the AABC *L. fasciola* brood (Figure 3), although additional analyses of its RADseq dataset are needed to resolve that issue.

There are well-known cases of a single genetic locus controlling a mimic polymorphism in other systems. In butterflies, polymorphic mimicry in wing pigmentation is controlled by an introgressed mimicry supergene in *Heliconius* species (Jay et al., 2018) and by mimicry alleles of the transcription factor *doublesex* (*dsx*) in some *Papilio* species (Palmer & Kronforst, 2020).

Note, however, that the *Lampsilis fasciola* mantle lure mimicry polymorphism differs in important ways from these butterfly systems. It is more complex because it involves putative models (darters and leeches) from disparate phyla rather than from similar morphospecies (other butterflies), thereby requiring polymorphic trait differentiation in pigmentation **and** in morphology (Figure 1b, c). It is also a case of aggressive mimicry (Jamie, 2017), different from the Müllerian mimicry of *Heliconius* (Kronforst & Papa, 2015) or the Batesian mimicry of *Papilio* (Kunte, 2009).

Persistence of *Lampsilis fasciola* mantle lure polymorphism across a broad geographic scale (Michigan to Alabama, Figure 2) is notable although the mechanism responsible for widespread maintenace is unclear. One hypothesized mechanism for the persistence of polymorphisms in a species or population is frequency-dependent selection, where selection for rare phenotypes are selected cause the ratio of phenotypes to vary over time (Ayala & Campbell, 1974). Frequency-dependent selection has been observed in other polymorphic mimicry systems (Shine, Brown & Goiran, 2022) and it has been suggested as a possible mechanism for persistence of the *L. fasciola* polymorphism (Zanatta, Fraley & Murphy, 2007; Barnhart, Haag & Roston, 2008; Hewitt, Haponski & Foighil, 2021). One criterion for frequency-dependent selection is that phenotype ratios oscillate over time as initially rare phenotypes become more successful. However, the historical (1954-1962) and contemporary (2017) data from Sharon Mills County Park (Figure 5) did not show evidence of such oscillation: the frequency of the more common “darter” lure – 84.2% vs. 85.2% - and the ratio of “darter” to “leech” lures – 48:9 vs. 23:4 - remained essentially the same for both time windows, although we of course lack data for intervening years. Theoretically, there are other mechanisms for balancing selection to maintain polymorphisms over long time-scales, including heterozygote advantage or opposing selection pressures favoring different alleles at polymorphic loci (Prout, 2000; Mérot et al., 2020), but underlying genetics of the *L. fasciola* polymorphism is unknown at this time and more data are clearly needed.

An integrative graphic outline of the *Lampsilis fasciola* mimetic system at the River Raisin study site was assembled containing 4 “darter” and 1 “leech mantle lure phenotypes together with their putative model species (Figure 7). The four putative River Raisin darter model species – *Etheostoma blennioides*, *E. exile*, *E. microperca* and *Percina maculata* – are all common and widespread members of the drainage’s ichthyofauna with 300-900 specimens of each species recovered from 30-100 sampling locations by the Smith et al. (1981) ecological survey. The relative uniformity of the “leech” mantle lure phenotype in the River Raisin and throughout the *L. fasciola* range (Figure 6f-j) stands in sharp contrast with much higher local and range-wide variation shown by “darter” lures (Figure 6a-e). That phenotypic lure disparity mirrors the collective phenotypic variability of darters vs. *Macrobdella decora*; darters are the second-most diverse fish clade in North America, with ∼170 species (Warren & Burr, 1994; Stein & Morse, 2000). Another possibility is that at least some *L. fasciola* “darter-like” lures across the mussel’s range are composite mimics of visual elements from more than one member of their local darter fauna, but that remains to be established.

While the behavior of mantle lures in *Lampsilis* mussels has been documented and studied for many decades (Ortmann, 1911; Kramer, 1970; Haag & Warren, 1999), detailed analysis of lure undulation behavior is currently lacking, and the relative importance of behavior versus coloration and morphology is not well understood. The lure undulation for both *L. cardium* and *L. fasciola* starts about 2/3rds of the way to the “posterior” (“tailed”) side of the lure, and then travels forward toward the “eyespot”-bearing “anterior”. This is quite different from the oscilatory “S” shaped routine swimming movements used by many fishes (Liao, 2007; Smits, 2019), however, it shares some resemblence to the “C” start behavior that many fishes use as an escape mechanism (Witt, Wen & Lauder, 2015). The unusual motion of the mantle lures may therefore be mimicing an escape behavior to some extent to strike.

Although the L. fasciola behaviors differ significantly from those exhibited by L. cardium, there appears to be no behavioral polymorphism to distinguish the darter from leech lure phenotypes. Our putative model for River Raisin *L. cardium* mantle lures is a species of pelagic minnow, *Pimephales notatus* (Figure 7), whose swimming behavior and ecology differs markedly from that of darters (Burress et al., 2017). Darters are primarily benthic in habit, have lost or greatly reduced their swim bladder (Demski, Gerald & Popper, 1973; Zeyl et al., 2016), and they swim by “hopping” across the river bed in a manner that is much more erratic and intermittent than the pelagic swimming behavior of most minnows (Winn, 1958). This matches a general difference observed between *L. cardium* and *L. fasciola* lures: *L. cardium* moves faster and more regularly in a highly synchronized way, in contrast with the erratic, often left-right-unsynchronized movements of *L. fasciola* lures. The only major difference in lure behavior between the “darter” and “leech” lures of *L. fasciola* is a slightly slower rate exhibited by the “darter” lures (Fig. 9a). Both *L. fasciola* morphs have a similar erratic motion, despite the polymorphism putatively modeling taxa from disparate phyla. Leeches swim by a dorsoventeral bending wave moving from head to tail, caused by a central pattern generator (CPG) comprised of interneurons in the ventral nerve cord (Jordan, 1998). This swimming behavior is very different from the lure undulations observed in the leech-like *L. fasciola* lures. Despite small differences in overall lure beat frequency between “darter” and “leech” mimics, it seems as though the polymorphic mimicry is only skin deep. The rhythmic movements of lure undulations are likely caused by a CPG, but we currently have no information about how these motor patterns are activated or modulated (Marder et al., 2005).

The *Lampsilis fasciola* mantle lure aggressive mimicry polymorphism is complex, each morph containing correlated morphological and pigmentation distinctions, however, the polymorphism does not appear to be linked to any large differences in lure behavior, or in the CPG that controls lure behavior. Our discovery of discrete within-brood inheritance of the *L. fasciola* lure polymorphism is of particular interest because it implies potential control by a single genetic locus. There are a number of parallel cases in the recent literature, *e.g.,* in butterflies, the regulation of polymorphic mimicry in wing pigmentation also involves single genetic loci (Jay et al., 2018; Palmer & Kronforst, 2020)Timmermans et al. (2020). Timmermans et al. (2020) used SNP data from *Papilio dardanus* to discover a genomic inversion associated with its mimetic polymorphism, and this approach is likely also tractable for *Lampsilis fasciola* given the availability of polymorphic broods – we are currently raising an additional polymorphic brood at the AABC. Mantle lures are a key adaptive trait in Lampsiline evolution and diversification (Hewitt, Haponski & Foighil, 2021) and *Lampsilis fasciola* is a promisinga highly tractable model system to uncover the genetics of lure development and variation in unionoid mussel.

