## Supplementary Materials for "Aggressive mimicry lure polymorphisms in the parasitic mussel *Lampsilis fasciola* model fish or leech host prey and differ in morphology and pigmentation, but not in display behavior"

Supplementary Table 1: Museum ID numbers, Raw reads, total clusters, and total loci in assembly from the ddRAD sequencing are displayed for each genotyped sample of *Lampsilis fasciola* and of the outgroup taxa. Individual *Lampsilis fasciola* lure phenotype designation followed Zanatta et al. (2007).

| Sample Name | Museum ID | | | Lure Phenotype | Raw reads | Total clusters | Average clustering depth | Loci in assembly |
|---|---|---|---|---|---|---|---|---|
| L_fasciola_AL_brood_1 | 306443 | 306443-1 | 306444-1 | Worm-like | 258664 | 97681 | 2.14 | 483 |
| L_fasciola_AL_brood_2 | 306444 | 306443-2 | 306443-1 | Darter-like | 5201836 | 1120710 | 3.28 | 25686 |
| L_fasciola_AL_brood_3 | 306445 | 306443-3 | 306444-2 | Worm-like | 5492519 | 1126749 | 3.4 | 25703 |
| L_fasciola_AL_brood_4 | 306446 | 306443-4 | 306443-2 | Darter-like | 2429494 | 632254 | 2.84 | 21398 |
| L_fasciola_AL_brood_5 | 306447 | 306443-5 | 306444-3 | Worm-like | 3152003 | 760260 | 3.02 | 23761 |
| L_fasciola_AL_brood_6 | 306448 | 306443-6 | 306443-3 | Darter-like | 3212851 | 810898 | 2.87 | 23434 |
| L_fasciola_AL_brood_7 | 306449 | 306443-7 | 306443-4 | Darter-like | 3649891 | 593765 | 4.22 | 25363 |
| L_fasciola_AL_brood_8 | 306450 | 306443-8 | 306443-5 | Darter-like | 4869307 | 1462723 | 2.29 | 19089 |
| L_fasciola_AL_brood_9 | 306451 | 306443-9 | 306444-4 | Worm-like | 3158818 | 718169 | 3.08 | 23033 |
| L_fasciola_AL_brood_10 | 306452 | 306443-10 | 306443-6 | Darter-like | 4000321 | 915881 | 3.12 | 24916 |
| L_fasciola_AL_brood_11 | 306453 | 306443-11 | 306444-5 | Worm-like | 5679854 | 1171842 | 3.35 | 25770 |
| L_fasciola_AL_brood_12 | 306454 | 306443-12 | 306443-7 | Darter-like | 4212783 | 979265 | 3.04 | 24693 |
| L_fasciola_AL_brood_13 | 306455 | 306443-13 | 306444-6 | Worm-like | 1300563 | 399134 | 2.51 | 12145 |
| L_fasciola_AL_brood_14 | 306456 | 306443-14 | 306443-8 | Darter-like | 4100372 | 1043360 | 2.79 | 23521 |
| L_fasciola_AL_brood_15 | 306457 | 306443-15 | 306443-9 | Darter-like | 5804293 | 1412102 | 2.91 | 25570 |
| L_fasciola_AL_brood_16 | 306458 | 306443-16 | 306444-7 | Worm-like | 1555906 | 427061 | 2.7 | 14099 |
| L_fasciola_AL_brood_17 | 306459 | 306443-17 | 306443-10 | Darter-like | 2073968 | 598680 | 2.59 | 13668 |
| L_fasciola_AL_brood_18 | 306460 | 306443-18 | 306444-8 | Worm-like | 6919783 | 1574429 | 3.08 | 25811 |
| L_fasciola_AL_brood_19 | 306461 | 306443-19 | 306443-11 | Darter-like | 3434210 | 829507 | 2.94 | 23708 |
| L_fasciola_AL_brood_20 | 306462 | 306443-20 | 306443-12 | Darter-like | 4778853 | 994416 | 3.35 | 25500 |
| L_fasciola_AL_brood_21 | 306463 | 306443-21 | 306444-9 | Worm-like | 2462560 | 590095 | 2.91 | 20588 |
| L_fasciola_AL_brood_22 | 306464 | 306443-22 | 306444-10 | Worm-like | 6600876 | 1406451 | 3.26 | 26080 |
| L_fasciola_AL_brood_23 | 306465 | 306443-23 | 306443-13 | Darter-like | 7090859 | 1628965 | 3.06 | 25932 |

| | | | | | | | | |
|---|---|---|---|---|---|---|---|---|
| L_fasciola_AL_brood_24 | 306466 | 306443-24 | 306444-11 | Worm-like | 4546435 | 1061394 | 3 | 24174 |
| L_fasciola_AL_brood_25 | 306467 | 306443-25 | 306444-12 | Worm-like | 5379577 | 1135906 | 3.35 | 25703 |
| L_fasciola_AL_brood_26 | 306468 | 306443-26 | 306444-13 | Worm-like | 5592652 | 1501130 | 2.67 | 23965 |
| L_fasciola_AL_brood_27 | 306469 | 306443-27 | 306444-14 | Worm-like | 4893957 | 825855 | 4.09 | 25924 |
| L_fasciola_AL_brood_28 | 306470 | 306443-28 | 306443-14 | Darter-like | 2596873 | 519103 | 3.59 | 22103 |
| L_fasciola_AL_brood_29 | 306471 | 306443-29 | 306443-15 | Darter-like | 3401334 | 883485 | 2.87 | 21377 |
| L_fasciola_AL_brood_30 | 306472 | 306443-30 | 306444-15 | Worm-like | 3876395 | 1014133 | 2.8 | 22072 |
| L_fasciola_AL_brood_31 | 306473 | 306443-31 | 306444-16 | Worm-like | 5391442 | 1246528 | 3.07 | 25009 |
| L_fasciola_AL_brood_32 | 306474 | 306443-32 | 306443-16 | Darter-like | 4365005 | 1084596 | 2.85 | 23030 |
| L_fasciola_AL_brood_33 | 306475 | 306443-33 | 306443-17 | Darter-like | 5116507 | 1117916 | 3.16 | 24667 |
| L_fasciola_AL_brood_34 | 306476 | 306443-34 | 306443-18 | Darter-like | 7480755 | 1601100 | 3.19 | 26163 |
| L_fasciola_AL_brood_35 | 306477 | 306443-35 | 306443-19 | Darter-like | 8121426 | 1825135 | 3.02 | 25972 |
| L_fasciola_AL_brood_36 | 306478 | 306443-36 | 306443-20 | Darter-like | 5521997 | 1414238 | 2.78 | 24163 |
| L_fasciola_AL_brood_37 | 306479 | 306443-37 | 306443-21 | Darter-like | 6562641 | 1579514 | 2.88 | 25476 |
| L_fasciola_AL_brood_38 | 306480 | 306443-38 | 306443-22 | Darter-like | 6303766 | 1596624 | 2.76 | 24448 |
| L_fasciola_AL_brood_39 | 306481 | 306443-39 | 306443-23 | Darter-like | 6206795 | 1488925 | 2.91 | 24648 |
| L_fasciola_AL_brood_40 | 306482 | 306443-40 | 306443-24 | Darter-like | 8630897 | 1891164 | 3.11 | 26176 |
| L_fasciola_AL_brood_41 | 306483 | 306443-41 | 306443-25 | Darter-like | 7293683 | 1716571 | 2.95 | 25604 |
| L_fasciola_AL_brood_42 | 306484 | 306443-42 | 306443-26 | Darter-like | 4896252 | 1193262 | 2.88 | 22829 |
| L_fasciola_AL_brood_43 | 306485 | 306443-43 | 306443-27 | Darter-like | 6098052 | 1471714 | 2.9 | 25074 |
| L_fasciola_AL_brood_44 | 306486 | 306443-44 | 306443-28 | Darter-like | 7495994 | 1698871 | 3.04 | 25701 |
| L_fasciola_AL_brood_45 | 306487 | 306443-45 | 306443-29 | Darter-like | 3937758 | 670698 | 4.06 | 24947 |
| L_fasciola_AL_brood_46 | 306488 | 306443-46 | 306443-30 | Darter-like | 6370942 | 1343655 | 3.26 | 25855 |
| L_fasciola_AL_brood_47 | 306489 | 306443-47 | 306443-31 | Darter-like | 5542864 | 1318463 | 2.96 | 24550 |
| L_fasciola_AL_brood_48 | 306490 | 306443-48 | 306443-32 | Darter-like | 6313913 | 1469606 | 2.98 | 24983 |
| L_fasciola_AL_brood_49 | 306491 | 306443-49 | 306443-33 | Darter-like | 3163000 | 789239 | 2.9 | 24776 |
| L_fasciola_AL_brood_50 | 306492 | 306443-50 | 306443-34 | Darter-like | 1728370 | 548529 | 2.35 | 17837 |
| L_fasciola_Huron_5 | 306493 | 306444-1 | 306445-1 | Darter-like | 953302 | 259898 | 2.8 | 10996 |
| L_fasciola_Huron_6 | 306494 | 306444-2 | 306446-1 | Worm-like | 1682931 | 362706 | 3.31 | 16809 |

| | | | | | | | | |
|---|---|---|---|---|---|---|---|---|
| L_fasciola_Huron_7 | 306495 | 306444-3 | 306446-2 | Worm-like | 746944 | 157212 | 3.29 | 10644 |
| L_fasciola_Huron_8 | 306496 | 306444-4 | 306446-3 | Worm-like | 1899689 | 402515 | 3.25 | 16584 |
| L_fasciola_Huron_9 | 306497 | 306444-5 | 306445-2 | Darter-like | 1213655 | 293090 | 2.97 | 11818 |
| L_fasciola_Huron_10 | 306498 | 306444-6 | 306445-3 | Darter-like | 7775910 | 1275602 | 3.87 | 22035 |
| L_fasciola_Huron_11 | 306499 | 306444-7 | 306445-4 | Darter-like | 1533281 | 295767 | 3.55 | 15386 |
| L_fasciola_NC_1 | 306500 | 306445-1 | 306447-1 | Darter-like | 1308813 | 254002 | 3.61 | 11873 |
| L_fasciola_NC_2 | 306501 | 306445-2 | 306447-2 | Darter-like | 4862573 | 852380 | 3.77 | 18321 |
| L_fasciola_NC_3 | 306502 | 306445-3 | 306447-3 | Darter-like | 663874 | 165869 | 2.95 | 9960 |
| L_fasciola_NC_4 | 306503 | 306445-4 | 306447-4 | Darter-like | 2610453 | 465228 | 3.76 | 13790 |
| L_fasciola_NC_5 | 306504 | 306445-5 | 306447-5 | Darter-like | 6927947 | 1459334 | 3.05 | 20804 |
| L_fasciola_NC_6 | 306505 | 306445-6 | 306447-6 | Darter-like | 1051195 | 202171 | 3.27 | 12415 |
| L_fasciola_NC_7 | 306506 | 306445-7 | 306447-7 | Darter-like | 1948092 | 382878 | 3.61 | 17101 |
| L_fasciola_NC_8 | 306507 | 306445-8 | 306447-8 | Darter-like | 3475751 | 669278 | 3.69 | 20683 |
| L_fasciola_NC_9 | 306508 | 306445-9 | 306447-9 | Darter-like | 5693936 | 1634946 | 2.46 | 22325 |
| L_fasciola_NC_10 | 306509 | 306445-10 | 306447-10 | Darter-like | 2175381 | 464794 | 3.38 | 17094 |
| L_fasciola_NC_11 | 306510 | 306445-11 | 306447-11 | Darter-like | 2189933 | 516643 | 3.05 | 17580 |
| L_fasciola_Redo_1 | 306511 | 306446-1 | 306448-1 | Darter-like | 1455864 | 327622 | 2.62 | 13478 |
| L_fasciola_Redo_2 | 306512 | 306446-2 | 306448-2 | Darter-like | 1839020 | 436418 | 2.43 | 13181 |
| L_fasciola_Raisin_2 | 306513 | 306447-1 | 306449-1 | Darter-like | 8235827 | 1716137 | 3.29 | 25555 |
| L_fasciola_Raisin_3 | 306514 | 306447-2 | 306449-2 | Darter-like | 6032935 | 1488448 | 2.85 | 25006 |
| L_fasciola_Raisin_4 | 306515 | 306447-3 | 306449-3 | Darter-like | 12947164 | 3587458 | 2.45 | 25245 |
| L_fasciola_Raisin_1 | 306516 | 306447-4 | 306449-4 | Darter-like | 6639384 | 1086218 | 3.97 | 23458 |
| L_fasciola_Raisin_5 | 306517 | 306447-5 | 306449-5 | Darter-like | 10059843 | 1997619 | 3.41 | 25363 |
| L_fasciola_Raisin_6 | 306518 | 306447-6 | 306449-6 | Darter-like | 8019689 | 1847955 | 3.01 | 25769 |
| L_fasciola_Raisin_7 | 306519 | 306447-7 | 306449-7 | Darter-like | 3816242 | 681697 | 3.95 | 24606 |
| L_fasciola_Raisin_8 | 306520 | 306447-8 | 306449-8 | Darter-like | 6117037 | 1282299 | 3.27 | 22439 |
| L_fasciola_Raisin_9 | 306521 | 306447-9 | 306450-1 | Worm-like | 5170380 | 775979 | 4.64 | 25798 |
| L_fasciola_Raisin_10 | 306522 | 306447-10 | 306449-9 | Darter-like | 761451 | 176858 | 3.14 | 11477 |
| L_fasciola_Raisin_11 | 306523 | 306447-11 | 306450-2 | Worm-like | 7140657 | 1670143 | 2.97 | 25519 |

| | | | | | | | | |
|---|---|---|---|---|---|---|---|---|
| L_fasciola_Raisin_12 | 306524 | 306447-12 | 306449-10 | Darter-like | 890521 | 203114 | 2.91 | 10582 |
| L_fasciola_Raisin_13 | 306525 | 306447-13 | 306449-11 | Darter-like | 1071361 | 225030 | 3.47 | 13512 |
| L_fasciola_Raisin_14 | 306526 | 306447-14 | 306449-12 | Darter-like | 3644379 | 946273 | 2.82 | 21995 |
| L_fasciola_Raisin_15 | 306527 | 306447-15 | 306449-13 | Darter-like | 3578043 | 482446 | 5.04 | 17514 |
| L_fasciola_Raisin_16 | 306528 | 306447-16 | 306449-14 | Darter-like | 2351544 | 114072 | 14.25 | 516 |
| L_fasciola_Raisin_17 | 306529 | 306447-17 | 306449-15 | Darter-like | 5272816 | 1304726 | 2.87 | 23305 |
| L_fasciola_Huron_1 | 306530 | 306448-1 | 306452 | Worm-like | 13366692 | 4050829 | 2.26 | 17555 |
| L_fasciola_Huron_2 | 306531 | 306448-2 | 306451-1 | Darter-like | 2819896 | 928226 | 2.24 | 20205 |
| L_fasciola_Huron_3 | 306532 | 306448-3 | 306451-2 | Darter-like | 662275 | 186602 | 2.66 | 7653 |
| L_fasciola_Huron_4 | 306533 | 306448-4 | 306451-3 | Darter-like | 4792093 | 855457 | 3.88 | 24512 |
| L_fasciola_AL_mom_1_10 | 306534 | 306449-1 | 306453-1 | Darter-like | 8095030 | 1840917 | 2.95 | 25420 |
| L_fasciola_AL_mom_2_21 | 306535 | 306449-2 | 306453-2 | Darter-like | 10329331 | 3504027 | 2.03 | 24488 |
| L_fasciola_AL_mom_3_16 | 306536 | 306449-3 | 306453-3 | Darter-like | 10384477 | 2987559 | 2.34 | 25056 |
| L_fasciola_Huron_12 | 306537 | 306450-1 | 306454-1 | Worm-like | 6906349 | 1672394 | 2.87 | 25281 |
| L_fasciola_Huron_13 | 306538 | 306450-2 | 306454-2 | Worm-like | 6955496 | 1670627 | 2.88 | 25593 |
| L_fasciola_Raisin_18 | 306539 | 306450-3 | 306454-3 | Worm-like | 5506215 | 1301878 | 3 | 25373 |
| L_fasciola_Raisin_19 | 306540 | 306451-1 | 306455-1 | Worm-like | 6611596 | 1524682 | 3.03 | 25604 |
| L_fasciola_Raisin_20 | 306541 | 306451-2 | 306455-2 | Worm-like | 4894495 | 1276608 | 2.74 | 24931 |
| L_fasciola_Raisin_21 | 306542 | 306451-3 | 306455-3 | Worm-like | 8396562 | 1736736 | 3.26 | 25490 |
| L_cardium_1 | 306543 | 306452-1 | 306456-1 | | 6864226 | 1710220 | 2.8 | 14625 |
| L_cardium_2 | 306544 | 306452-2 | 306456-2 | | 4898330 | 1091622 | 3.11 | 13433 |
| L_cardium_3 | 306545 | 306452-3 | 306456-3 | | 7109883 | 2005565 | 2.5 | 14563 |
| L_cardium_4 | 306546 | 306452-4 | 306456-4 | | 4637077 | 997208 | 3.27 | 13860 |
| S_nasuta_1 | 306547 | 306453 | 306457 | | 4544989 | 1169260 | 2.55 | 10441 |

Supplementary Table 2: Summary data on individual mantle lure display field recordings. Video recordings were taken during the summer of 2018 at Sharon Mills (Fig. 2a) and Hudson Mills (Fig. 2b). Average movement length and interval was calculated from frame number from a 120fps video recording using a Go Pro Hero 6.

| File | Average Movement duration (L) | Average Movement duration (R) | Average Interval (L) | Average Interval (R) | Interval Standard Deviation (L) | Interval Standard Deviation (R) | Proportion of Movements Synchronized | Lure Phenotyp e | Time | Temperature °C | Date | Site |
|---|---|---|---|---|---|---|---|---|---|---|---|---|
| GH010073 | 0.18 | 0.18 | 1.50 | 1.99 | 1.28 | 1.65 | 0.44 | "leech" | 11:26 | 16.4 | 7/10/18 | Hudson Mills |
| GH010074 | 0.14 | 0.14 | 1.72 | 1.69 | 1.73 | 1.73 | 0.56 | "leech" | 14:09 | 18.5 | 7/9/18 | Sharon Mills |
| GH010599 | 0.22 | 0.23 | 13.17 | 8.71 | 20.34 | 19.34 | 0.04 | "darter" | 11:29 | 18.3 | 7/10/18 | Hudson Mills |
| GH010601 | 0.21 | 0.22 | 1.02 | 1.25 | 0.52 | 0.33 | 0.48 | "leech" | NA | NA | 7/10/18 | Hudson Mills |
| GH010602 | 0.17 | 0.17 | 1.19 | 1.52 | 0.62 | 0.69 | 0.32 | "leech" | 13:36 | 20.6 | 7/10/18 | Hudson Mills |
| GH010603 | 0.23 | 0.21 | 1.20 | 1.14 | 1.22 | 1.10 | 0.24 | "darter" | 13:57 | 20.5 | 7/10/18 | Hudson Mills |
| GH010075 | 0.15 | 0.15 | 1.77 | 1.58 | 1.79 | 1.78 | 0.27 | "leech" | 2:48 | 18.2 | 7/9/18 | Sharon Mills |
| GH010598 | 0.27 | 0.27 | 1.80 | 1.75 | 1.19 | 1.17 | 0.80 | "darter" | 11:18 | 18.2 | 7/10/18 | Hudson Mills |
| GH010597 | 0.31 | 0.30 | 1.30 | 1.27 | 0.57 | 0.50 | 0.93 | "leech" | 11:06 | 18.1 | 7/10/18 | Hudson Mills |
| GH010595 | 0.22 | 0.22 | 1.51 | 1.99 | 1.17 | 1.04 | 0.42 | "leech" | 10:34 | 17.6 | 7/10/18 | Hudson Mills |
| GH010056 | 0.24 | 0.24 | 3.12 | 3.00 | 1.89 | 2.26 | 0.38 | "darter" | 12:59 | 21.3 | 6/11/18 | Sharon Mills |
| GH010077 | 0.19 | 0.22 | 1.83 | 1.66 | 1.91 | 1.76 | 0.18 | "darter" | 11:26 | 16.4 | 7/10/18 | Hudson Mills |
| GH010055 | 0.29 | 0.29 | 2.34 | 2.34 | 1.32 | 1.31 | 0.60 | "darter" | 11:59 | 21.8 | 6/11/18 | Sharon Mills |
| GH010016 | 0.35 | 0.37 | 1.83 | 1.81 | 0.50 | 0.47 | 0.53 | "darter" | NA | NA | 6/12/18 | Sharon Mills |
| GH010057 | 0.30 | 0.26 | 9.87 | 7.40 | 11.22 | 8.65 | 0.30 | "darter" | 13:16 | 21.4 | 6/11/18 | Sharon Mills |
| GH010593 | 0.15 | 0.15 | 0.99 | 0.88 | 0.58 | 0.59 | 0.39 | "leech" | 14:09 | 20.7 | 7/9/18 | Sharon Mills |
| GH010062 | 0.14 | 0.14 | 2.27 | 2.77 | 1.76 | 1.84 | 0.24 | "darter" | 10:21 | 19.2 | 7/4/18 | Hudson Mills |
| GH010064 | 0.17 | 0.18 | 1.10 | 1.10 | 0.42 | 0.42 | 0.98 | "leech" | NA | 20.8 | 7/4/18 | Hudson Mills |
| GH010065 | 0.12 | 0.13 | 1.21 | 1.66 | 0.92 | 0.83 | 0.45 | "darter" | 14:23 | 21 | 7/4/18 | Hudson Mills |
| GH010063 | 0.14 | 0.14 | 1.05 | 1.05 | 0.29 | 0.31 | 0.82 | "darter" | 11:13 | 19.6 | 7/4/18 | Hudson Mills |
| GH010579 | 0.12 | 0.12 | 0.88 | 1.29 | 0.55 | 0.47 | 0.26 | "leech" | 11:02 | 20 | 7/4/18 | Hudson Mills |
| GH010580 | 0.13 | 0.13 | 1.56 | 1.82 | 1.13 | 1.43 | 0.67 | "leech" | 11:52 | 21.3 | 7/4/18 | Hudson Mills |
| GH010581 | 0.18 | 0.18 | 2.17 | 2.51 | 5.16 | 5.67 | 0.54 | "leech" | 13:36 | 23.3 | 7/4/18 | Hudson Mills |
| GH010582 | 0.10 | 0.11 | 1.94 | 2.16 | 2.76 | 2.79 | 0.19 | "darter" | 14:18 | 23.4 | 7/4/18 | Hudson Mills |
| GH010583 | 0.11 | 0.12 | 0.99 | 1.12 | 0.91 | 0.82 | 0.60 | "darter" | 14:52 | NA | 7/4/18 | Hudson Mills |
| GH010068 | 0.12 | 0.11 | 2.79 | 2.73 | 2.71 | 2.37 | 0.41 | "darter" | NA | NA | 6/12/18 | Sharon Mills |
| GH010048 | 0.15 | 0.14 | 8.05 | 7.67 | 5.94 | 5.60 | 0.39 | "darter" | NA | NA | 6/7/18 | Sharon Mills |
| GH010060 | 0.17 | 0.17 | 0.28 | 0.28 | 0.01 | 0.01 | 1.00 | cardium | NA | NA | 6/5/20 | Sharon Mills |
| GH010163 | 0.23 | 0.23 | 0.55 | 0.55 | 0.12 | 0.12 | 1.00 | cardium | NA | NA | 6/5/20 | Sharon Mills |
| GH010620 | 0.25 | 0.25 | 0.48 | 0.48 | 0.05 | 0.05 | 1.00 | cardium | NA | NA | 5/31/21 | Sharon Mills |
| GH010618 | 0.25 | 0.25 | 0.41 | 0.41 | 0.10 | 0.10 | 1.00 | cardium | NA | NA | 6/1/21 | Sharon Mills |

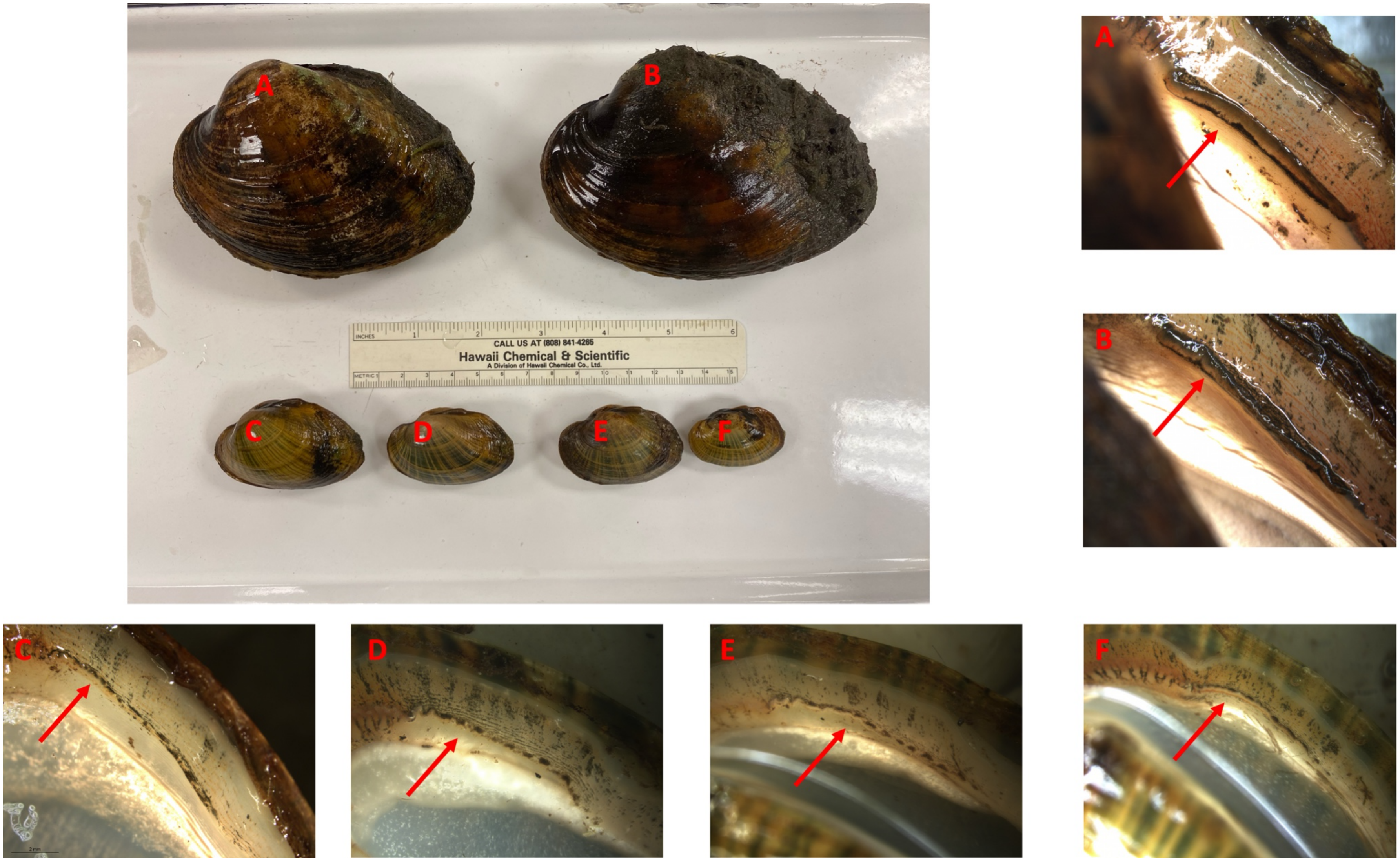


Supplementary Figure 1: A group photograph of 6 River Raisin Sharon Mills (MI) male lampsiline mussels - 2 *Lampsilis cardium* (A, B) and 4 *Lampsilis fasciola* (C-F) taken in May 2023 - together with individual photographs of their respective right mantle lure rudiments (arrows). All 4 *L. fasciola* mantle lures were "darter-like" with mottled coloration and minute marginal extensions.

### Group 1

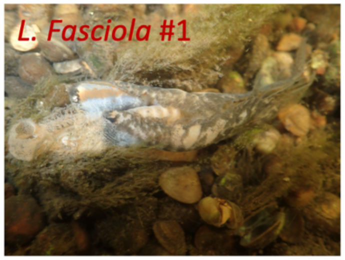


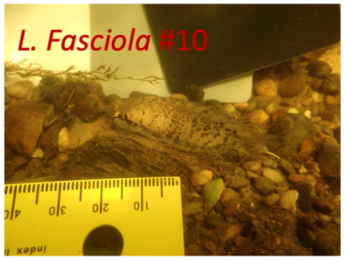


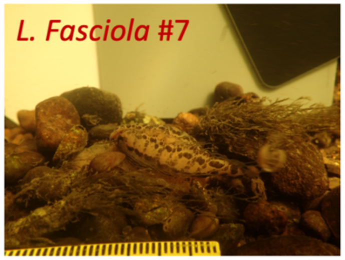


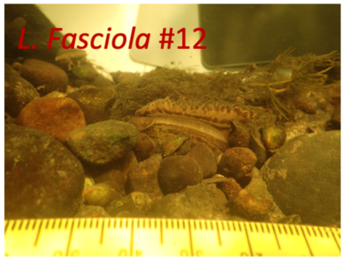


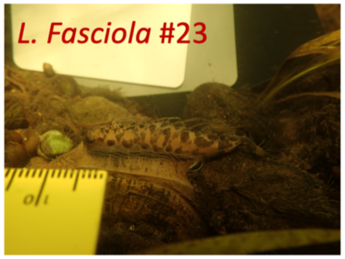


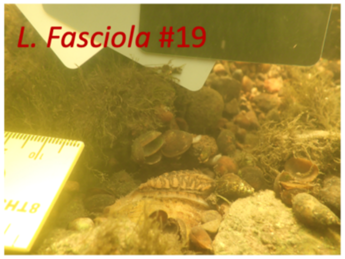


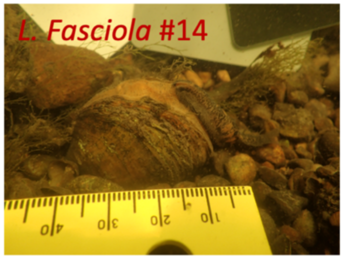


### Group 2

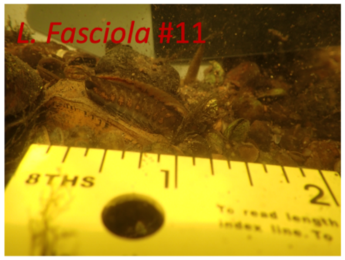


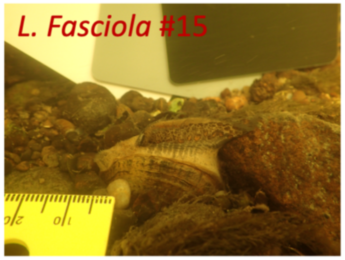


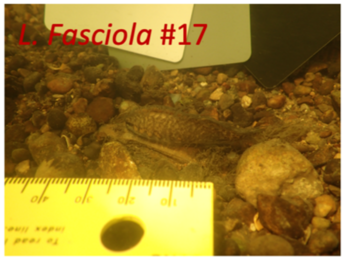

Group 3

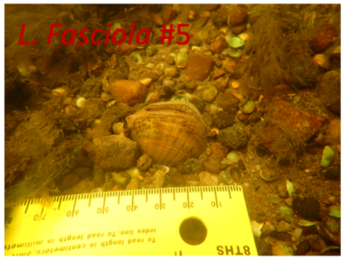
L. Fasciola #5

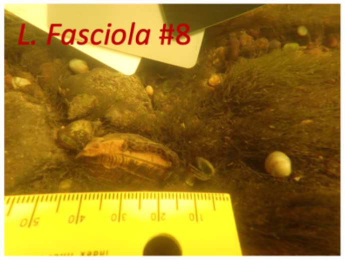
L. Fasciola #8

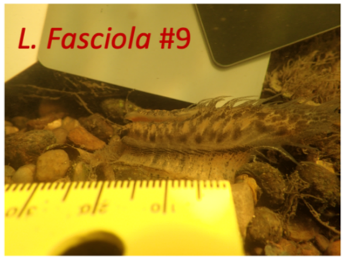
L. Fasciola #9

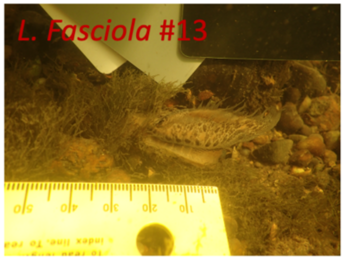
L. Fasciola #13

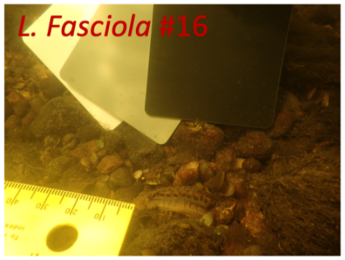
L. Fasciola #16

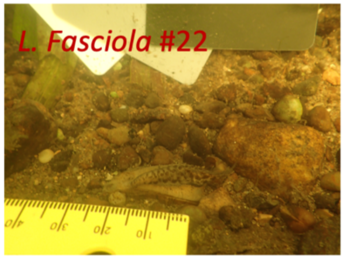
L. Fasciola #22

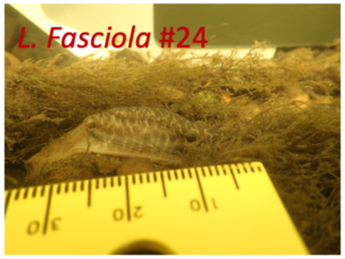
L. Fasciola #24

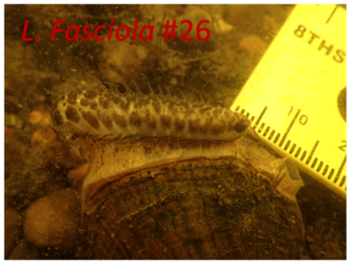
L. Fasciola #26
8THS

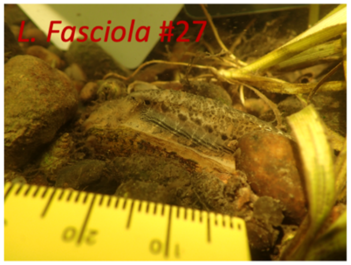
L. Fasciola #27

Group 4

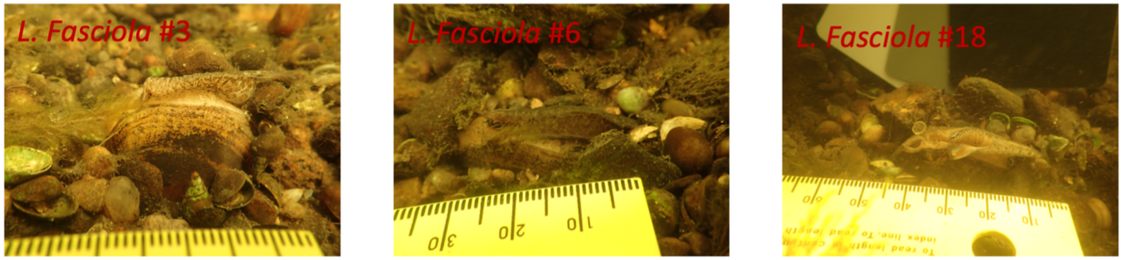


Group 5

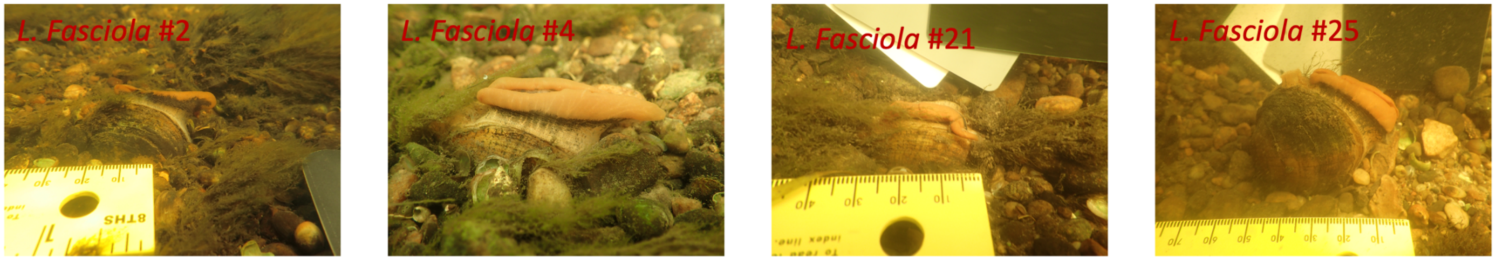


Supplementary Figure 2: Photographs from 27 *Lampsilis fasciola* lures taken at Sharon Mills (Fig. 2a) in Summer of 2017. Groups are defined by morphological similarity and individual numbers refer to order in which photographs were taken.

### Mussel brood notes

Date : 7-13-09

Species: L. fasciola

Source: Paint Rock R.

| ID number | Length (mm) | Width (mm) | Height (mm) | Mass (g) |
|---|---|---|---|---|
| LF01 | 54 | 26 | 38 | 32.32 |

* Bright orange and black mantle lure *

**A. Suspension volume (ml):** 250 (Adjust to give ~20 per drop)

| Drop # | Undeveloped eggs | Open glochidia | Closed glochidia | Open after salt |
|---|---|---|---|---|
| 1 | | 29 | Ø | 1 |
| 2 | | 28 | 1 | 1 |
| 3 | | 22 | 1 | 1 |
| 4 | | 20 | 1 | 1 |
| 5 | | 23 | 3 | Ø |
| 6 | | 17 | 1 | 1 |
| 7 | | 18 | Ø | Ø |
| 8 | | 29 | 3 | Ø |
| 9 | | 26 | 2 | Ø |
| 10 | | 26 | 4 | 1 |
| Sums | B | C 238 | D 16 | E 6 |

**Total glochidia = (C + D) * A/2 =** 31750

**Viable glochidia = (C - E) * A/2 =** 29000 912 viable glo.

7.25 L

29 REB

Supplementary Figure 3: Alabama Aquatic Biodiversity Center data sheet documenting the mantle lure phenotype (bracketed with red *s) and the larval brood size of the gravid Paint Rock River female *Lampsilis fasciola* used to establish the inaugural AABC cultured unionid brood in 2009.

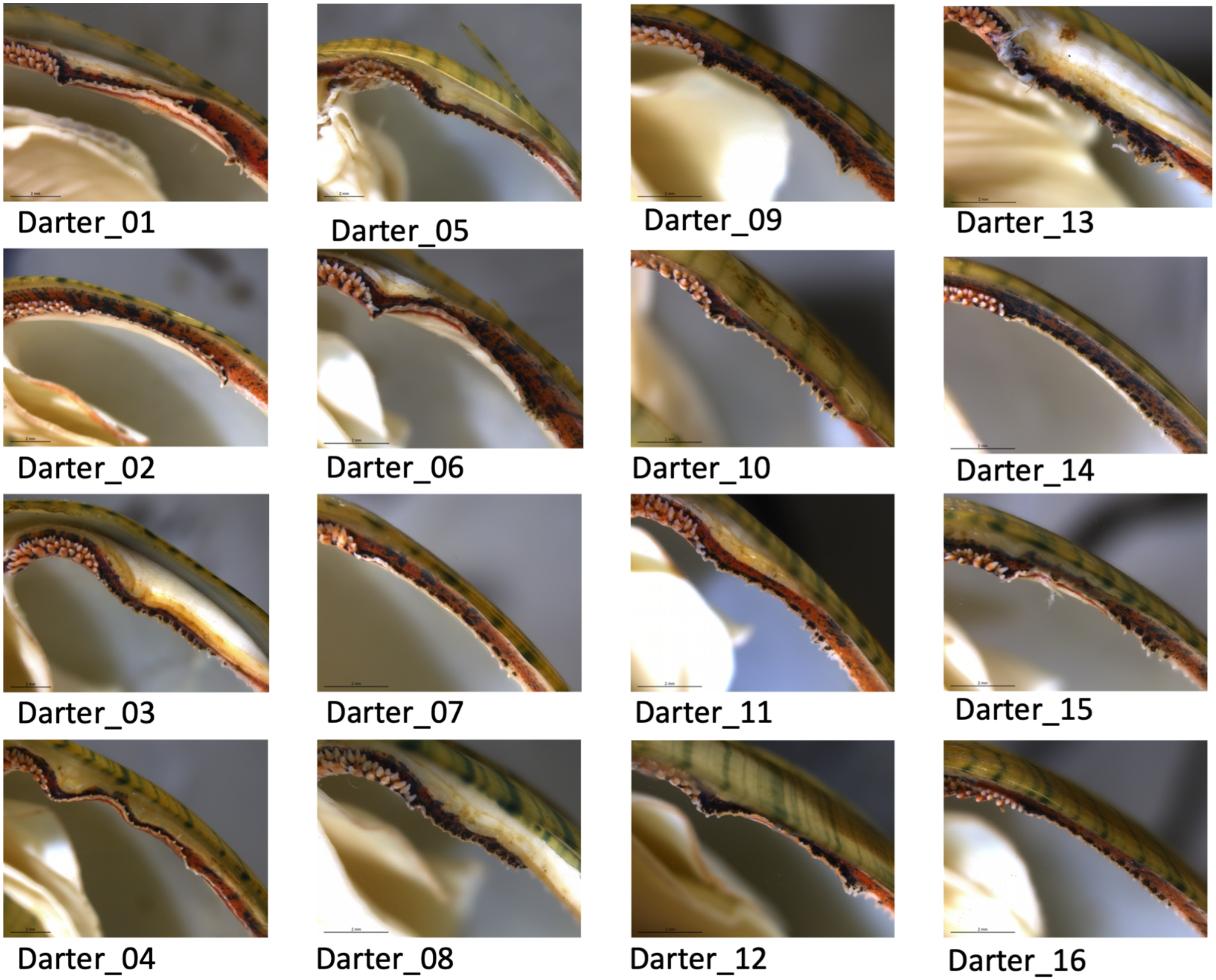
Darter_01
Darter_05
Darter_09
Darter_13
Darter_02
Darter_06
Darter_10
Darter_14
Darter_03
Darter_07
Darter_11
Darter_15
Darter_04
Darter_08
Darter_12
Darter_16

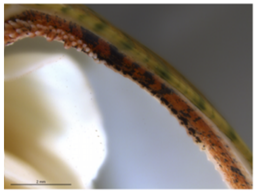
Darter_17

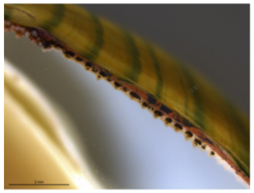
Darter_18

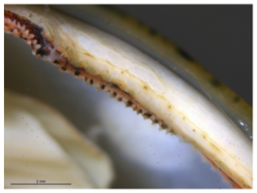
Darter_19

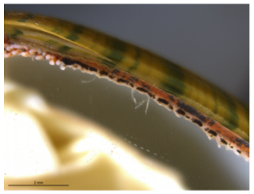
Darter_20

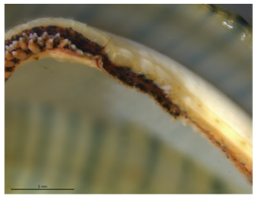
Darter_21

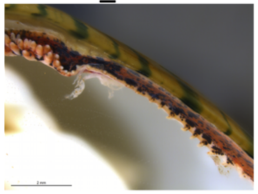
Darter_22

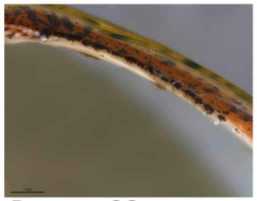
Darter_23

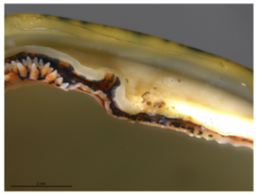
Darter_24

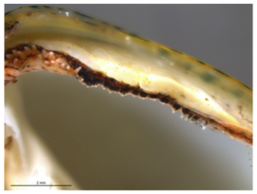
Darter_25

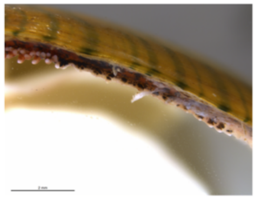
Darter_26

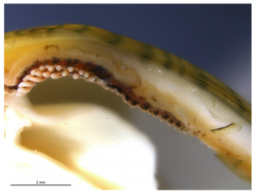
Darter_27

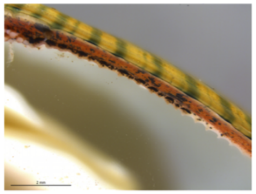
Darter_28

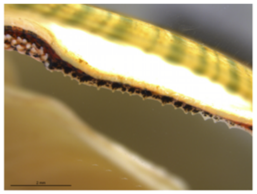
Darter_29

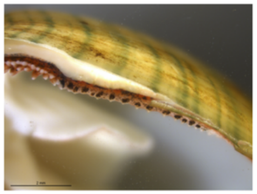
Darter_30

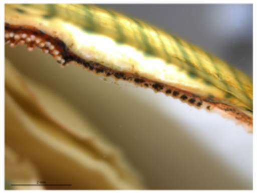
Darter_31

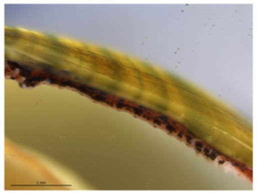
Darter_32

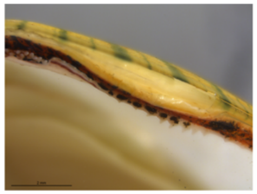
Darter_33

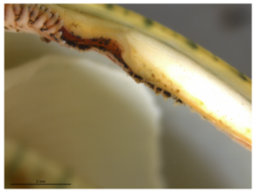
Darter_34

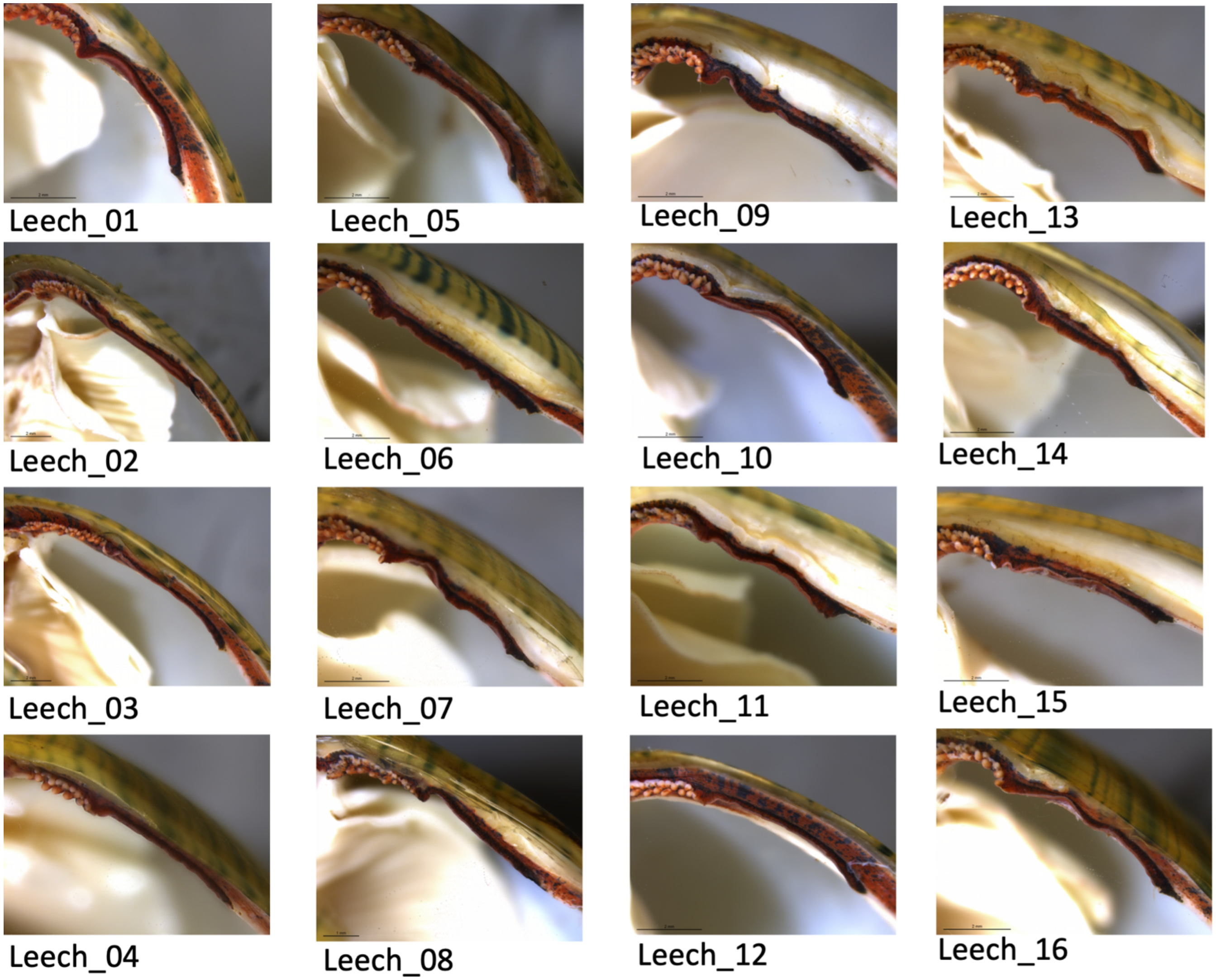


Supplementary Figure 4: Photographs of *Lampsilis fasciola* lure structure taken from 50 full or half-siblings raised from a single gravid female at the Alabama Aquatic Biodiversity Center in 2009. Each individual is categorized based on whether it has a darter-like or worm-like lure phenotype.

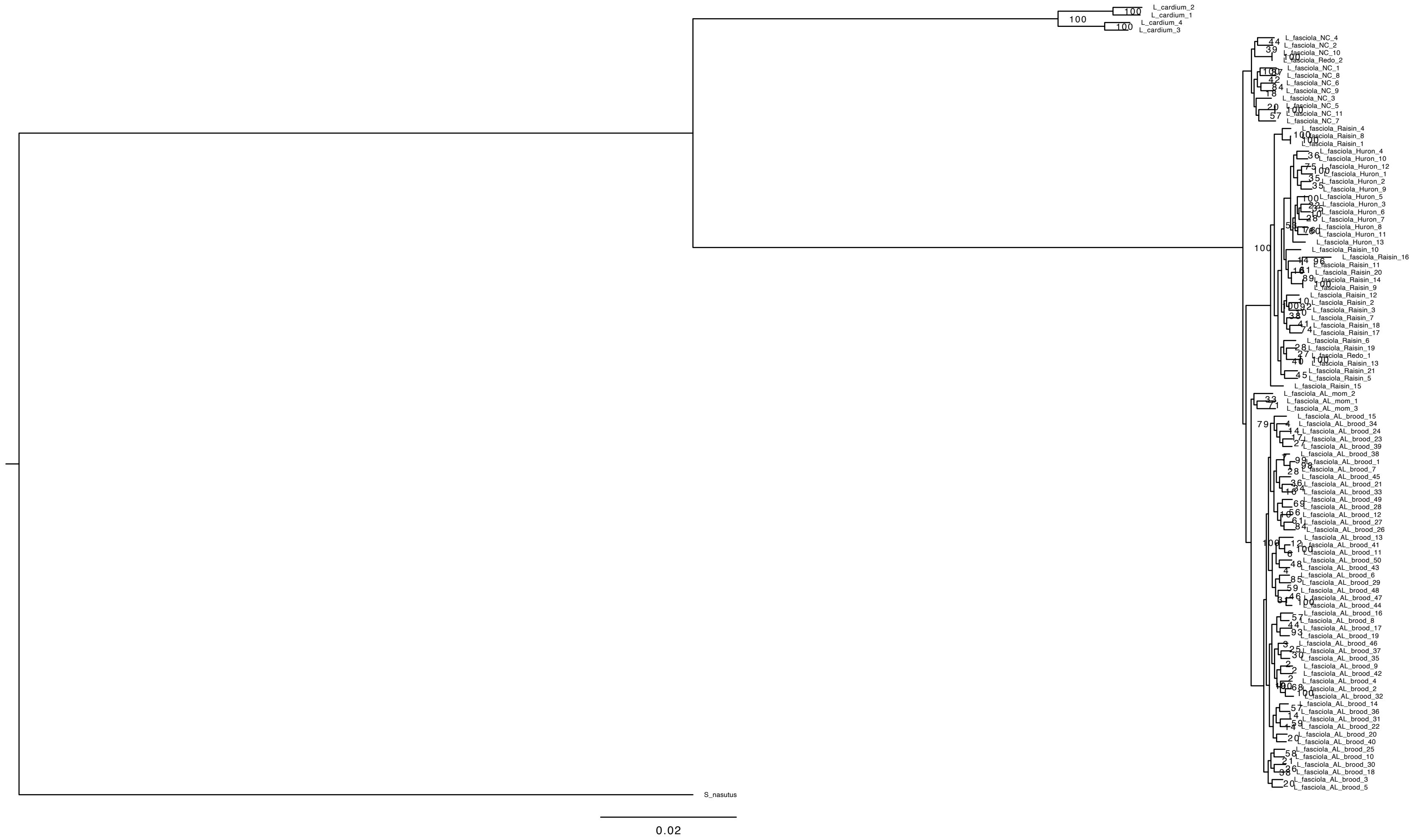


Supplementary Figure 5: Maximum likelihood phylogeny of *Lampsilis fasciola* mussels created with RAxML v8.2.8 using a general time reversible model. Support for each node was determined using 100 fast parametric bootstrap replications. Bootstrap values are adjacent to each node. Scale bar represents mean number of base pair substitutions per site.

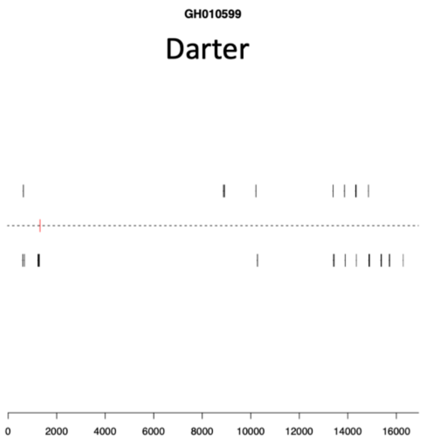
GH010599
Darter
0
2000
4000
6000
8000
10000
12000
14000
16000

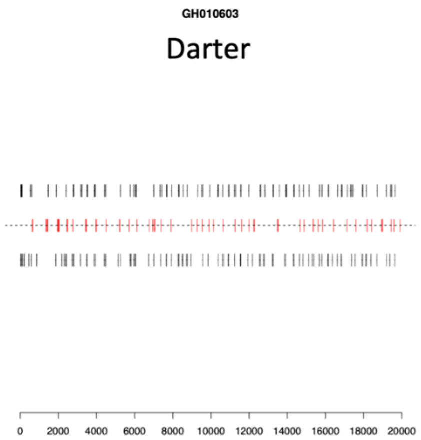
GH010603
Darter
0
2000
4000
6000
8000
10000
12000
14000
16000
18000
20000

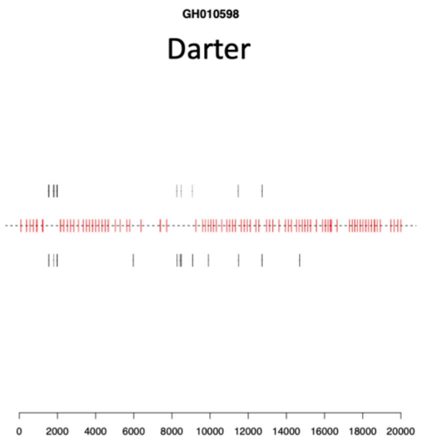
GH010598
Darter
0
2000
4000
6000
8000
10000
12000
14000
16000
18000
20000

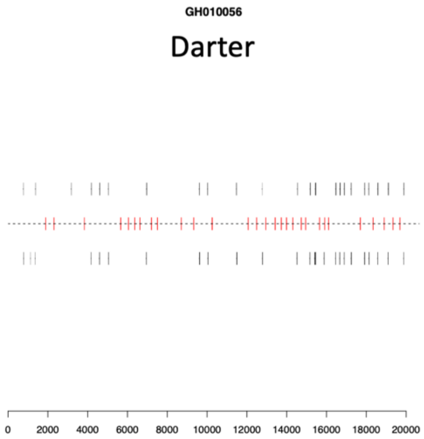
GH010056
Darter
0
2000
4000
6000
8000
10000
12000
14000
16000
18000
20000

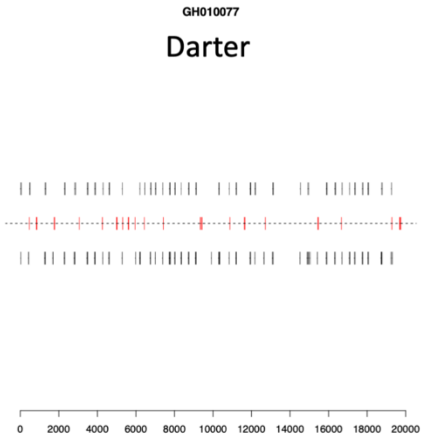
GH010077
Darter
0
2000
4000
6000
8000
10000
12000
14000
16000
18000
20000

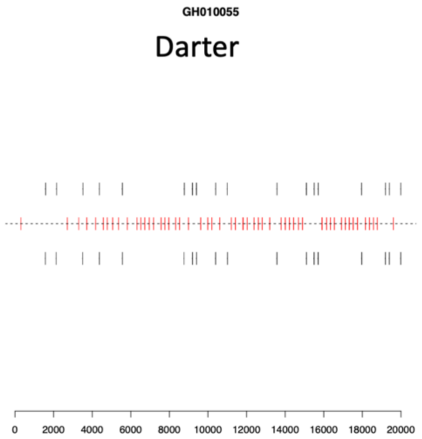
GH010055
Darter
0
2000
4000
6000
8000
10000
12000
14000
16000
18000
20000

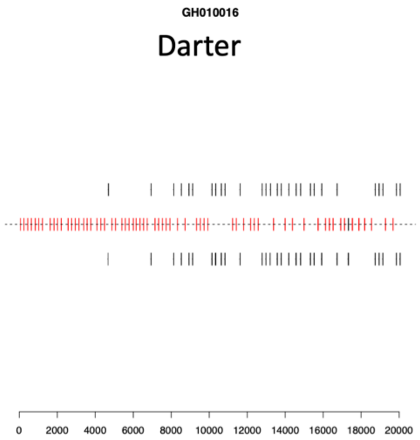
GH010016
Darter
0 2000 4000 6000 8000 10000 12000 14000 16000 18000 20000

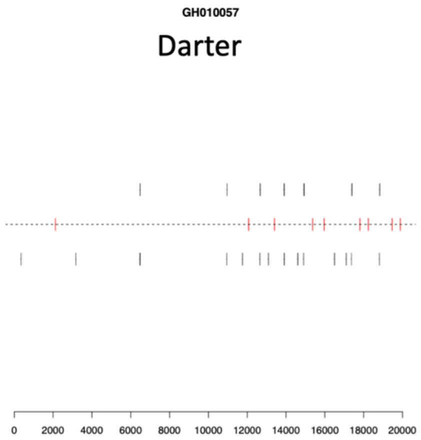
GH010057
Darter
0 2000 4000 6000 8000 10000 12000 14000 16000 18000 20000

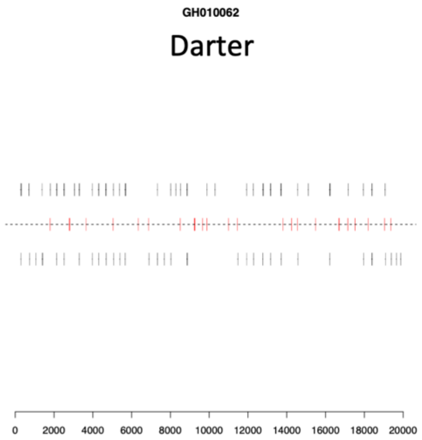
GH010062
Darter
0 2000 4000 6000 8000 10000 12000 14000 16000 18000 20000

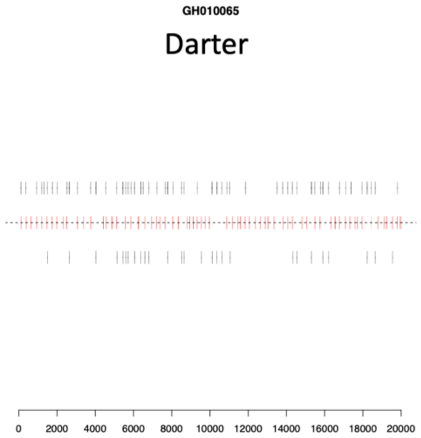
GH010065
Darter
0 2000 4000 6000 8000 10000 12000 14000 16000 18000 20000

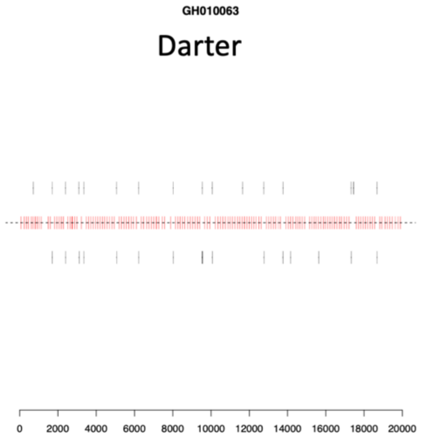
GH010063
Darter
0 2000 4000 6000 8000 10000 12000 14000 16000 18000 20000

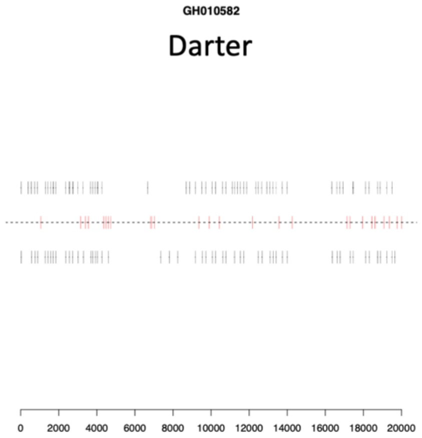
GH010582
Darter
0 2000 4000 6000 8000 10000 12000 14000 16000 18000 20000

GH010583

Darter

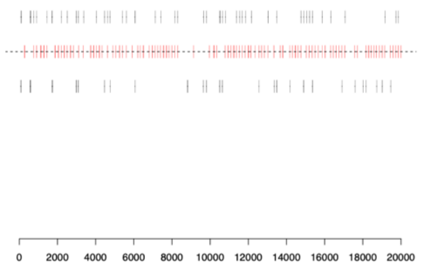


GH010068

Darter

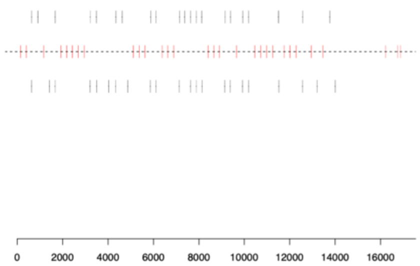


GH010048

Darter

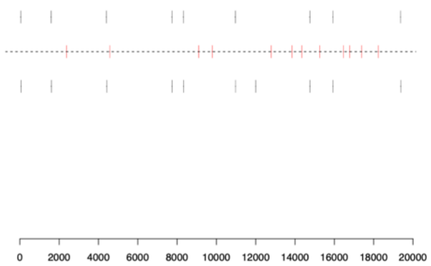

**GH010073**

### Leech

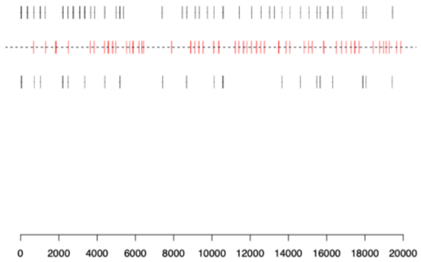


**GH010074**

### Leech

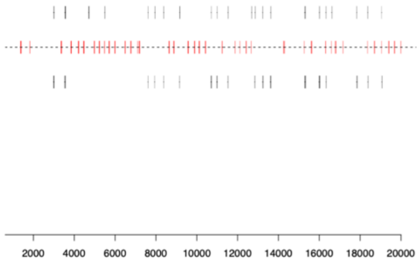


**GH010601**

### Leech

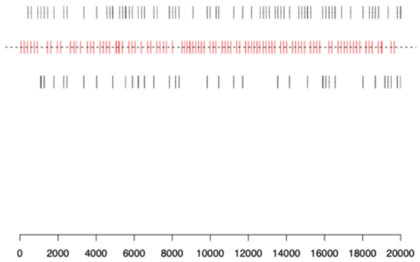


**GH010602**

### Leech

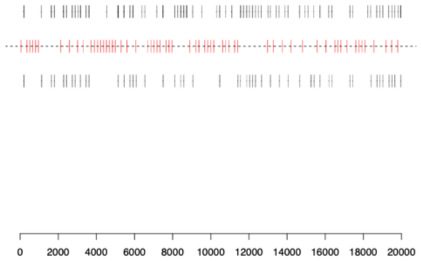


**GH010075**

### Leech

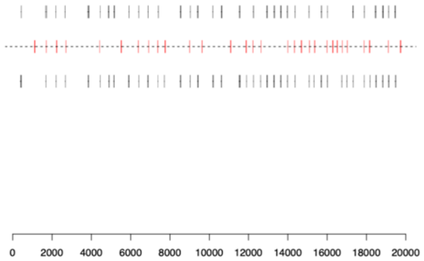


**GH010597**

### Leech

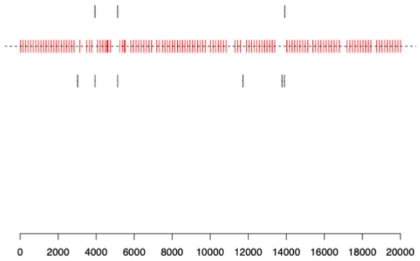

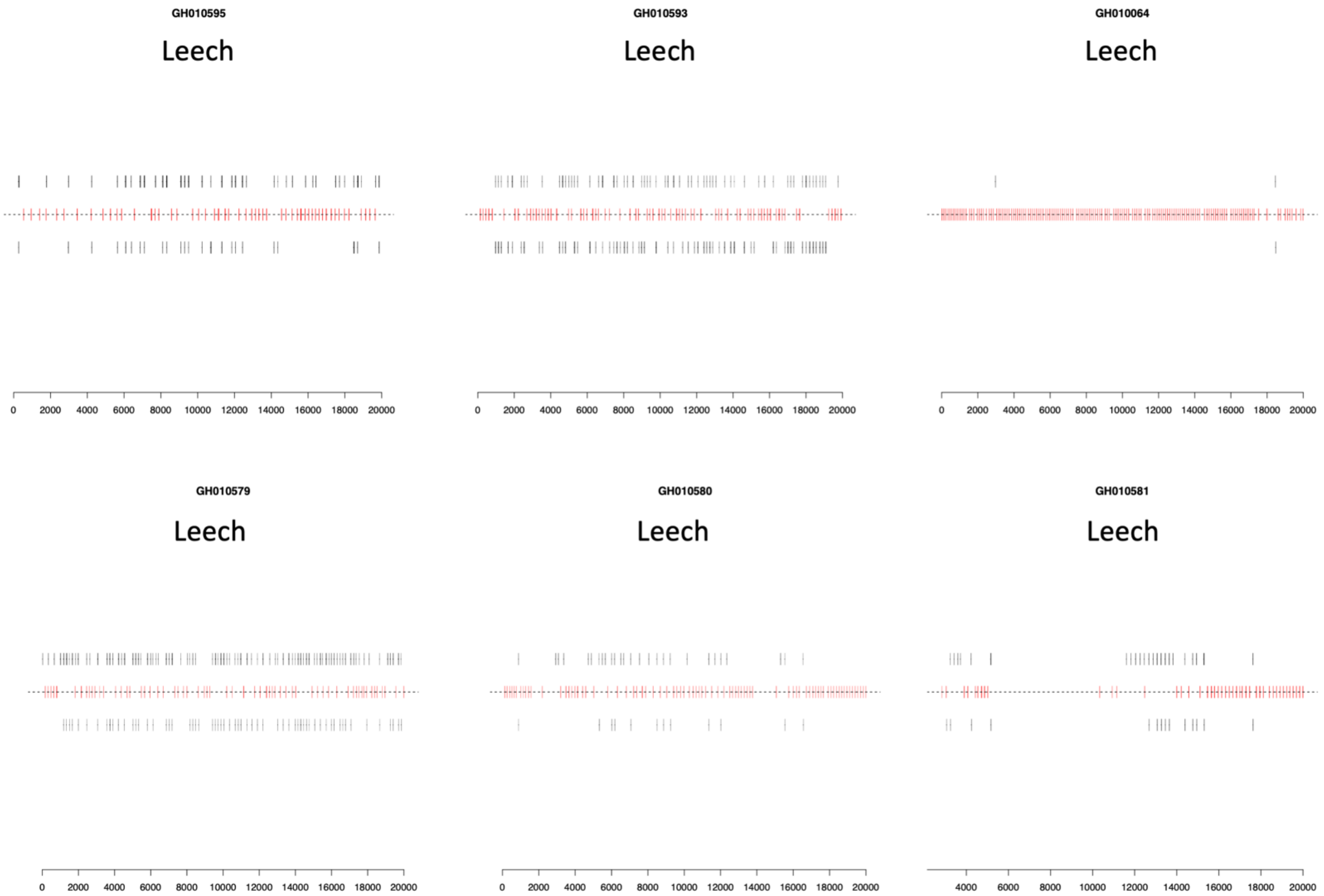
GH010595
Leech
0 2000 4000 6000 8000 10000 12000 14000 16000 18000 20000
GH010593
Leech
0 2000 4000 6000 8000 10000 12000 14000 16000 18000 20000
GH010064
Leech
0 2000 4000 6000 8000 10000 12000 14000 16000 18000 20000
GH010579
Leech
0 2000 4000 6000 8000 10000 12000 14000 16000 18000 20000
GH010580
Leech
0 2000 4000 6000 8000 10000 12000 14000 16000 18000 20000
GH010581
Leech
4000 6000 8000 10000 12000 14000 16000 18000 20000

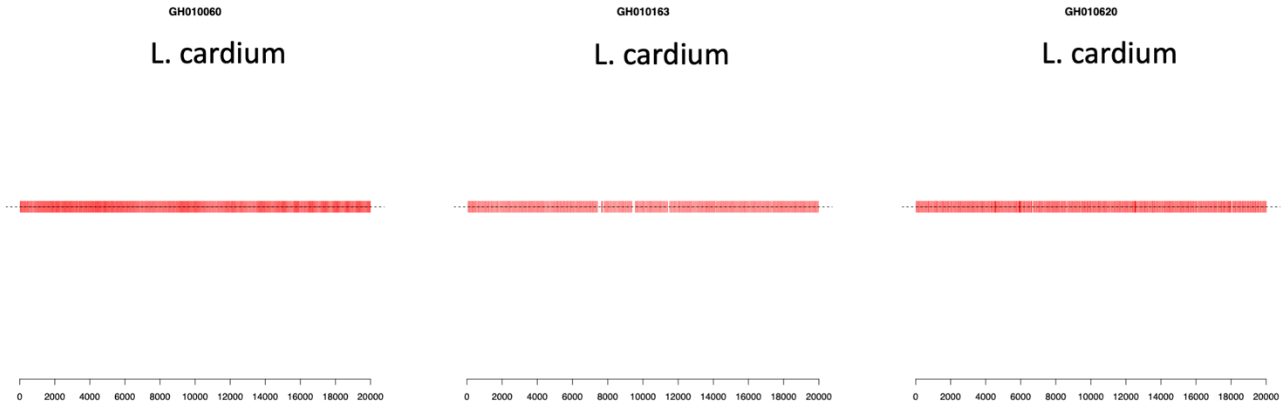


Supplementary Figure 6: Gait analyses for *Lampsilis fasciola* and *Lampsilis cardium* lure behavior. X axis denotes frame number. Videos were taken in Summer of 2018 from Sharon Mills (Fig. 2a) and Hudson Mills (Fig. 2b). Red lines on the center dotted line represent synchronized left/right movements, and black lines above the center line represent left side movements and below represent right side movements. Each graph is labelled as either a darter-like *L. fasciola*, leech-like *L. fasciola,* or *L. cardium.*

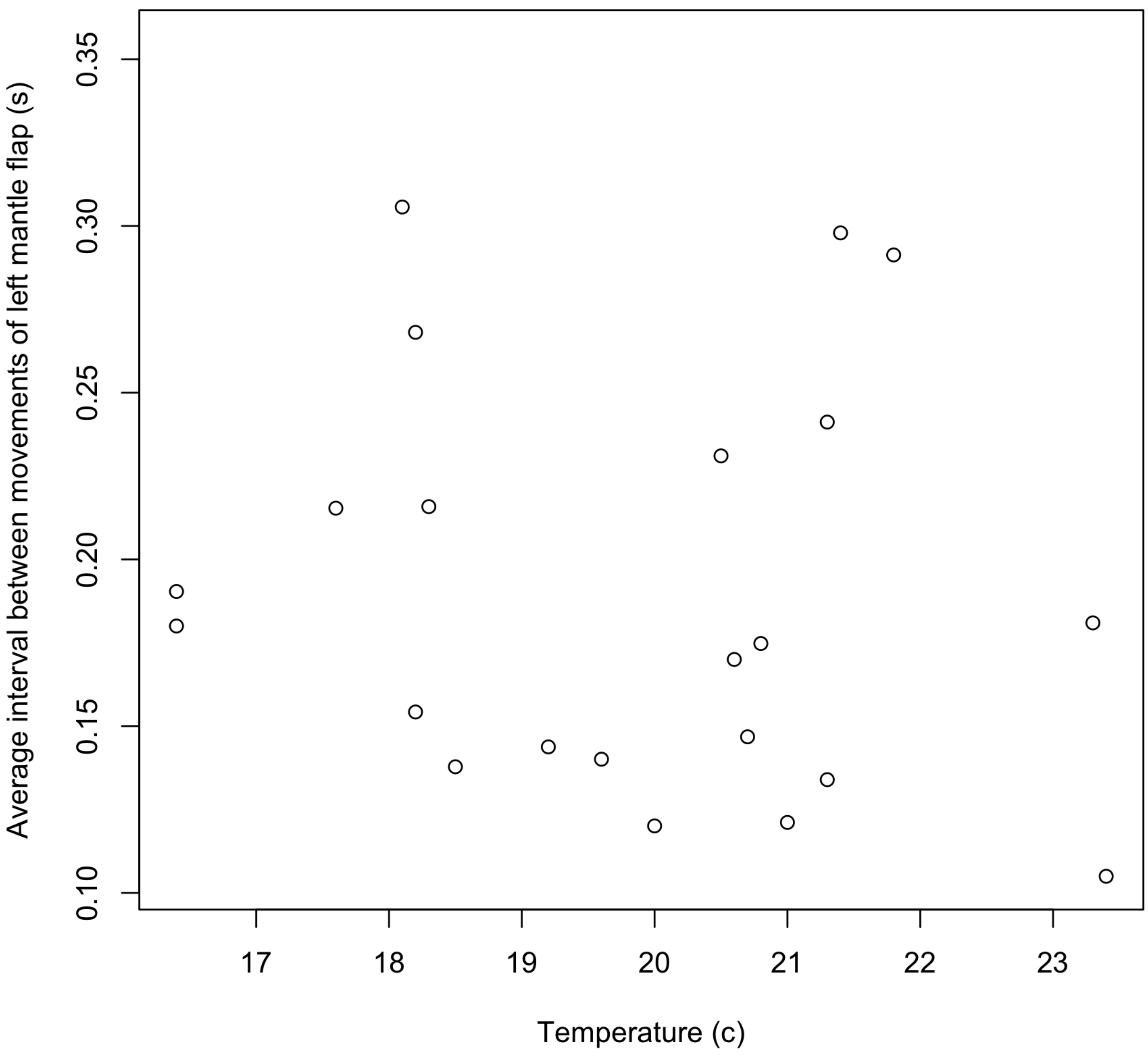


Supplementary Figure 7: Scatterplot showing the relationship between the average time interval between lure movements (left mantle flap) and temperature for *Lampsilis fasciola* and *Lampsilis cardium* lure displays videotaped in Summer 2018 and Sharon Mills (Fig. 2a) and Hudson Mills (Fig. 2b).
